# The eduWOSM: a benchtop advanced microscope for education and research

**DOI:** 10.1101/2024.08.10.607181

**Authors:** N.J. Carter, D.S. Martin, J.E. Molloy, R.A. Cross

## Abstract

To improve access to advanced optical microscopy in educational and resource-limited settings we have developed the eduWOSM (<u>edu</u>cational <u>W</u>arwick <u>O</u>pen <u>S</u>ource <u>Mic</u>roscope), an open hardware platform for transmitted-light and epifluorescence imaging in up to 4 colours, including single molecule imaging. EduWOSMs are robust, bright, compact, portable and ultra-stable. They are controlled entirely by open source hardware and software, with an option for remote control from a webpage. Here we describe the core eduWOSM technology and benchmark its performance using 3 example projects, single fluorophore tracking of tubulin heterodimers within gliding microtubules, 4D (deconvolution) imaging/tracking of chromosome motions in dividing human cells, and automated single particle tracking in vitro and in live cells with classification into subdiffusive, diffusive and superdiffusive motion.

## Introduction

Since its inception^1^, optical microscopy has been a central tool of scientific discovery, and of effective scientific training. In our own era, the open source movement is dramatically improving access to advanced microscopy^2^ driven by the advent of inexpensive cameras, light sources, microcontrollers/computers and a burgeoning global maker community that design and freely-share CAD and 3D-printable resources. Open source software is now of central importance in optical microscopy for generic image processing and microscope control (for example, ImageJ^3 4^, FIJI^5^, ICY^6^, μManager^7^,Pycro-Manager^8^), for data management (for example OMERO^9^) and, increasingly, for more computationally-intensive and specialised approaches, including adaptive data collection^10,11^. Open source hardware can in principle be equally important^12^ and a number of projects have already reached maturity^2,13^, including relatively complex STORM^14^ and light sheet^15^ microscopes and components^16^. The motivation for own project, the <u>Edu</u>cational <u>W</u>arwick <u>O</u>pen <u>S</u>ource <u>M</u>icroscope (eduWOSM) has been to create an optimised pladorm for high-resolution fluorescence microscopy in educational settings. We sought to design a modular, readily-reconfigurable pladorm with exceptional optical performance and mechanical stability that is driven by open-source, embedded-software electronics that allow instrument control via a local computer or remotely over TCP/IP. To maximise utility and lower the barriers to access, we wanted to minimise cost, and make the project fully open source. The novel features of the eduWOSM (***Fig. 1***) include a CNC-machined chassis that gives outstanding mechanical and thermal stability, an integrated, purpose-designed microcontroller with both digital and analogue lines, a miniaturised 4-colour LED light engine and an ergonomic, mouse-sized keypad and scroll wheel for manual control. Power comes from an integral, small form-factor, standard ATX PC power supply. Images are recorded using an inexpensive CMOS camera connected by fast USB3 data link to a PC running μManager.

**Figure 1.**
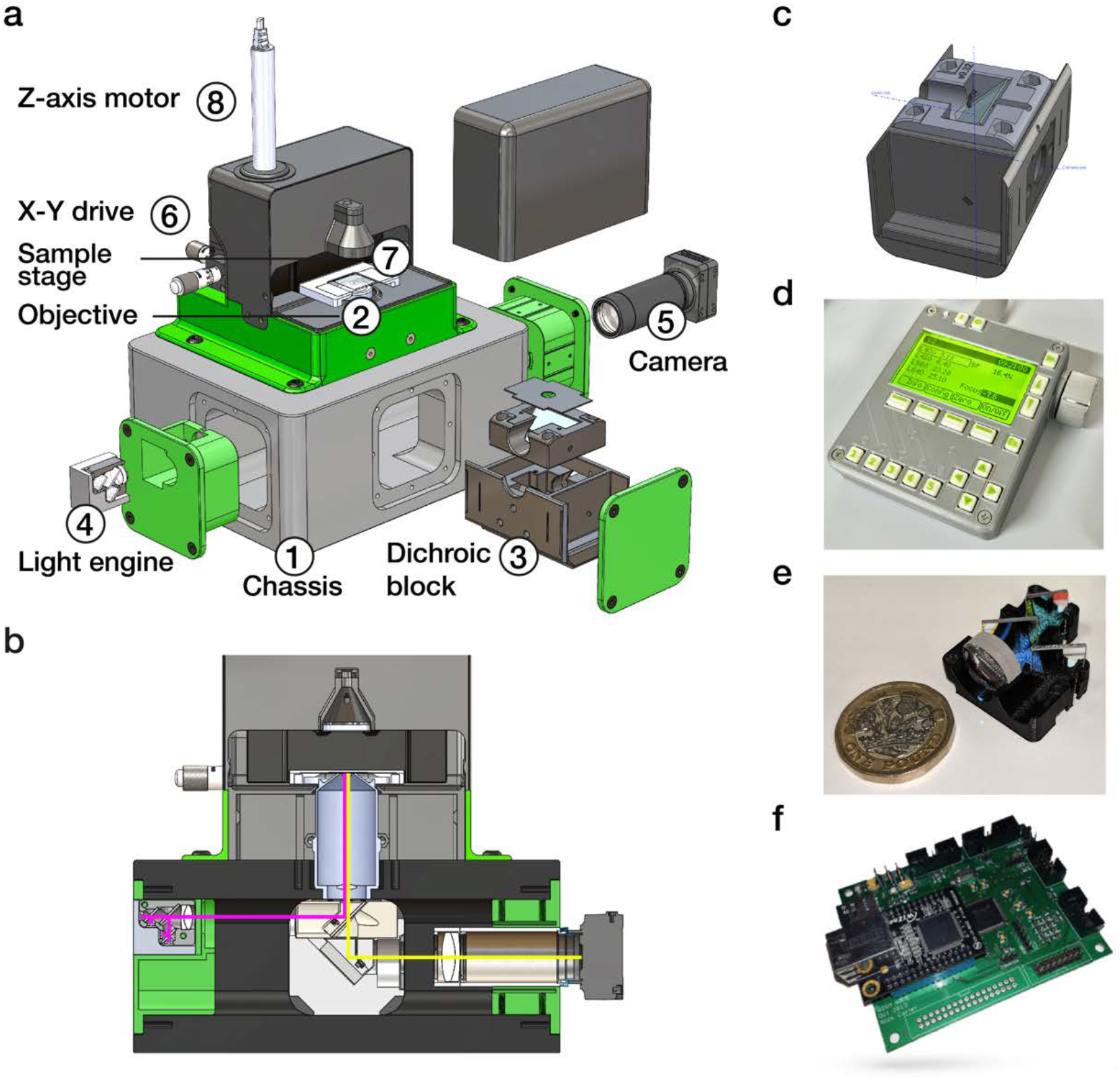
The eduWOSM. **a**| The eduWOSM chassis (1) is CNC machined from solid aluminium. A 100x NA 1.3 oil immersion objective (2) screws directly into the block and becomes monolithic with it. The four faces of the block carry identical ports, leading to a common central space beneath the objective. The dichroic block (3) slides into one of the ports and locks in position beneath the objective. The 4-colour LED light engine (4) and the camera (5) fit into 2 other ports. A 3-axis micrometer stage (6) is fixed to the top of the block and carries the sample holder (7). Slides are mounted to the sample holder using a pair of magnets. Z-axis motion of the stage is via a stepper motor (8). **b**| section through the microscope showing the optical path. Excitation path in magenta, emission path in yellow. **c**| Dichroic block. **d**| Keypad **e**| Light engine **f**| Microcontroller board.

Here, we report our designs for the core eduWOSM technologies, and benchmark their performance in 3 example experiments (***Figs 2-4***), all of which have recently been performed by first year undergraduate students on Warwick’s MSci Integrated Natural Sciences course^17^.

**Figure 2.**
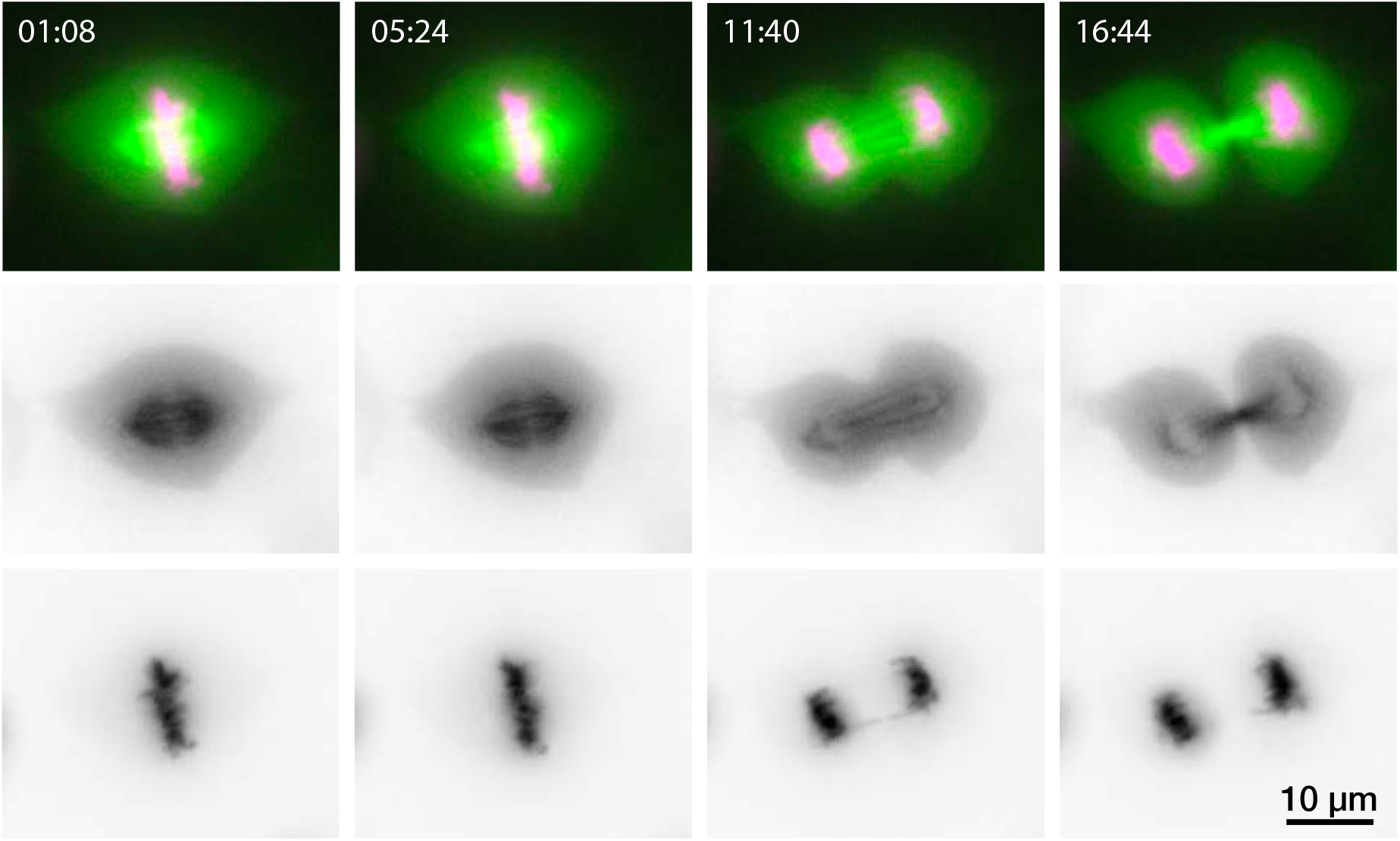
HeLa cell mitosis. Live imaging of a mitotic HeLa cell with a standard-configuration eduWOSM (100x objective). Representative video frames showing (left to right) mitotic spindle at metaphase, early anaphase, late anaphase and telophase. The top row shows combined tubulin (green) and chromatin (magenta) channels. The rows beneath show the two channels separately at each time point, in reverse contrast for clarity. The full recording consists of 400 frames taken at 4 sec intervals (***Supp. Video 1***).

**Figure 3.**
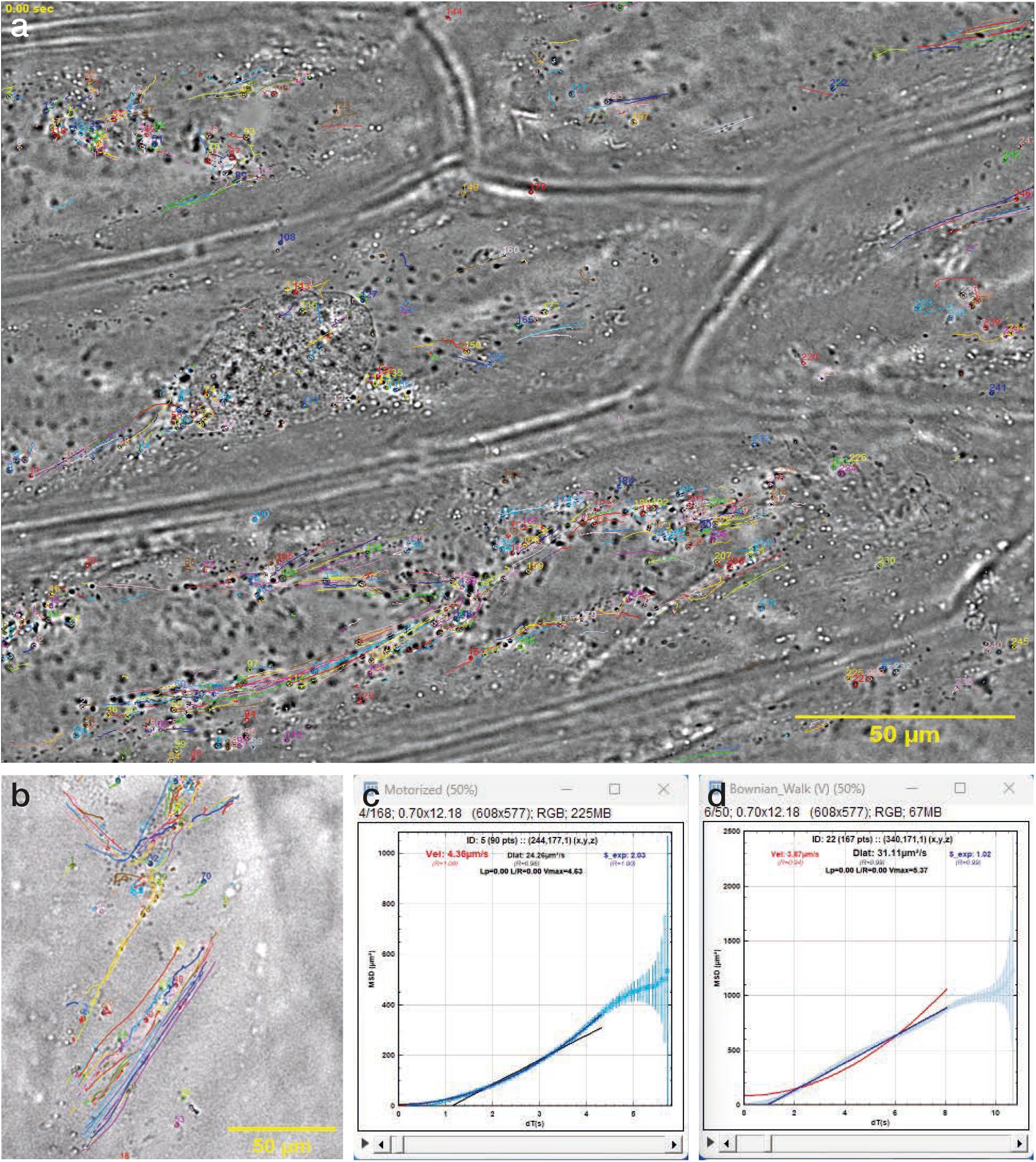
Quantifying diffusion in living cells. **a**| Imaging of a layer of living onion cells using the eduWOSM transmission illumination. Note the large field obtained at the diffraction-limited resolution of the 100x objective. The trajectories of 200 particles, obtained by automated tracking, are marked. Please zoom in for detail. **b**| Still from example movie (***Supp. Video 2***). **c**,**d**| The automated WOSM tracker macro (***Supp. File 1***) calculates MSD (mean squared displacement) for all pairs of position-time values along each trajectory and fits expressions for diffusion, subdiffusion and superdiffusion, returning a diffusion constant and an R (goodness-of-fit) value for each. **c**| example of a superdiffusive trajectory. **d**| example of a purely diffusive trajectory.

**Figure 4.**
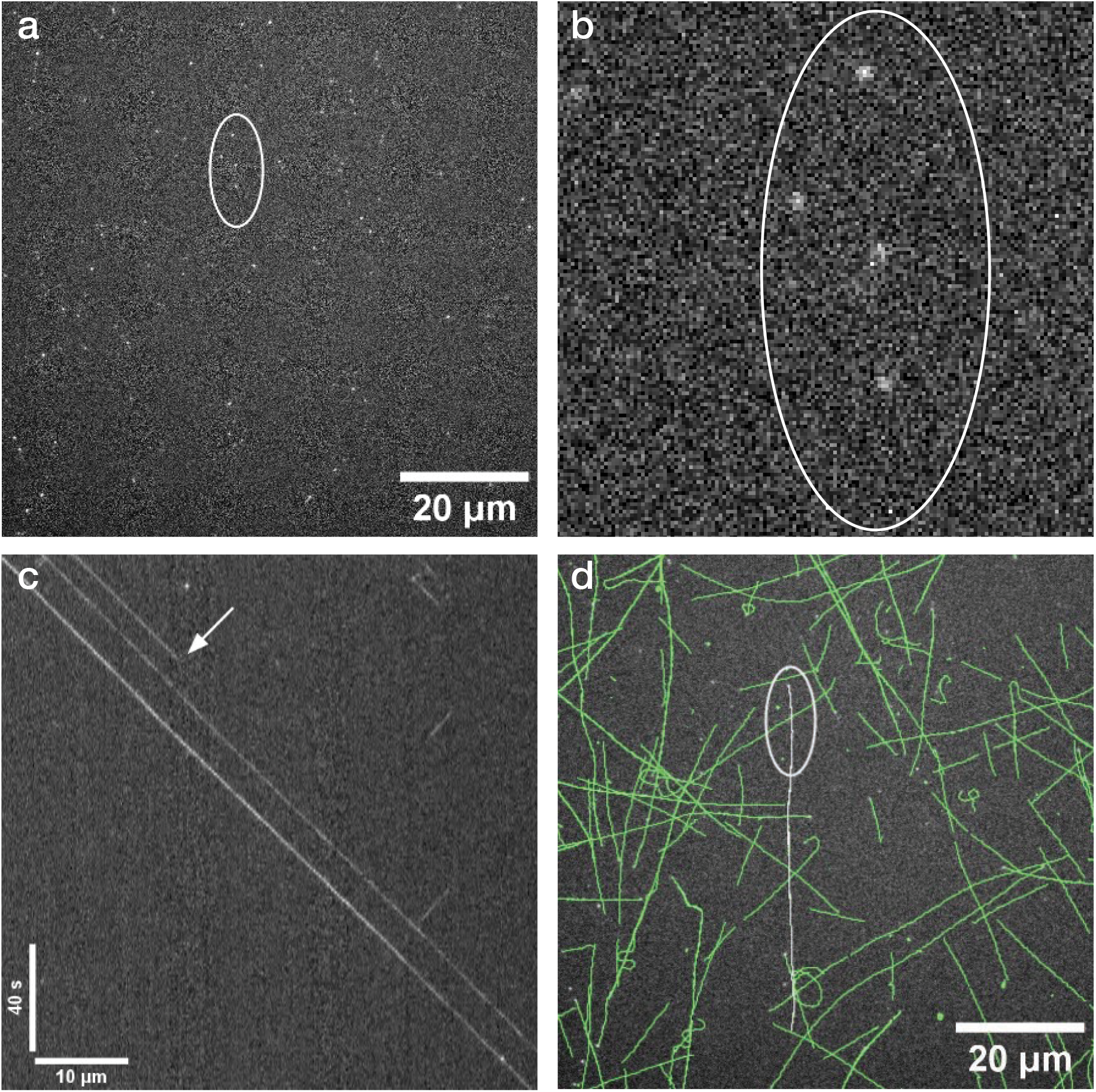
Tracking single fluorophores. Demonstration of single molecule imaging capabilities of the eduWOSM using gliding microtubules, sparsely labelled with HiLyte 647. **a**| Snapshot of microtubules. Spots corresponds to single HiLyte 647 fluorophores, illuminated by the red LED, exposure time 500 ms. **b**| Pixel-level view of the region marked. **c**| Kymograph showing that fluorophores on the marked microtubule glide at a constant velocity of 530 nm/s and that single-step photobleaching occurs (white arrow), indicating that the eduWOSM is successfully tracking single fluorophores. **d**| Trajectories of all microtubules in the field, constructed by superposing single fluorophore trajectories (***Supp. Video 3***).

## Results and Discussion

The eduWOSM **chassis** is a CNC-machined aluminium block (***Fig. 1***). An oil immersion objective (Nikon 100x 1.3NA Plan Fluor) screws directly into the chassis and becomes monolithic with it, giving outstanding short- and long-term mechanical and thermal stability. Each side of the block hosts an identical port. The ports accept 3D printed plastic sledges carrying componentry. Inserting components into the ports automatically aligns them along a common optical axis. The 3-axis **sample stage** mounts to the top of the block via pre-drilled metric-threaded holes. Mechanical stability over 1 hour is appreciably better than any of our commercial microscope systems.

The **eduWOSM light engine** is a miniaturised 4-colour LED illuminator (***Fig. 1***) that delivers illumination in UV, red, green and blue. Light from the 4 high-power LEDs is combined using dichroics and a projector lens that are both housed within the light engine. The 4 colours emerge aligned to a common optical axis. Once set during construction, the alignment is stable. Power to the LEDs (and other devices) is supplied by the eduWOSM power board, controlled by the **eduWOSM microcontroller** (below), giving 16-bit LED current control (0-700 mA) and μs switching, allowing for example precisely-timed strobed illumination.

Light from the light engine is deflected into the objective back aperture by the **dichroic block** (***Fig. 1***). The block is 3D printed in two halves. Its upper element is a multi-bandpass filter that reflects each of the 4 illuminating colours but transmits the fluorescence they generate to a second (high-quality, lambda/10) broadband mirror mounted directly below. Light striking this second mirror is reflected into a tube lens that illuminates the camera sensor. We were initially concerned that a plastic dichroic block would be unstable, but we find that housing it within the massive aluminium eduWOSM optical block confers great stability.

The **camera** (***Fig. 1***) is a CMOS camera with single fluorophore sensitivity (∼70% QE), 2448×2048 pixels and 12-bit resolution. A USB3.1 interface enables full-frame video capture at up to 35 frames per second. Image ROI cropping and/or pixel binning can be used to increase frame rate and decrease noise. By keeping the light path short we minimise optical leverage, improving image stability.

For imaging multi-colour fluorescence we collect the colours in sequence and synchronize light engine illumination intervals to the camera frame exposure start signal (available as a configurable TTL pulse on the camera auxiliary output). Captured frames can be overlaid in real-time within μManager so that users see a live multi-colour image. Our approach gives diffraction-limited resolution over a flat-field ∼230 μm across (92 nm pixels). Our use of a flat-field (plan) achromatic rather than apochromatic objective saves money and gives a brighter image, because there are fewer lens elements.

By default, samples are sandwiched between 2 coverslips and clipped to the **sample holder** (***Fig. 1***) using magnets. X and Y translation is done manually using micrometer drives chosen for their mechanical stability. Z-axis (focus) motion is via a stepper motor, allowing 100 nm steps. After mounting a specimen, a plastic hood can be snapped on magnetically to exclude room light and reduce convection and acoustic noise at the sample. An LED **transmission illuminator** is incorporated into the hood. The stage and sample cover also accommodate traditional sample slides. For temperature and gas control, the entire eduWOSM can be placed in an incubator.

Both local and remote control of the microscope are enabled by the **eduWOSM microcontroller** (***Fig. 1***). Input/output functionality includes high-speed 16-bit digital to analogue outputs, configurable digital I/O lines, stepper/servo motor control via optional modules and PWM or analogue outputs. There is appreciable spare capacity. Firmware updates for the controller are downloaded from WOSM.net website and are installed on the integrated micro-SD card. The micro-SD also stores user-defined scripts. Control of the microcontroller is via μManager, from a web browser, via telnet, via scripts stored locally on the micro-SD card or via the **eduWOSM keypad**.

The **eduWOSM keypad** is 3D printed and fractionally larger than a computer mouse. It enables control of the microcontroller using physical keys and a very precise rotary encoder. The keypad LCD screen reports current settings. The rotary encoder changes its function according to the screen mode setting, allowing it for example to control focus position or LED power.

The **user experience** is straightforward. On boot, the eduWOSM microcontroller negotiates an IP address using DHCP and then sets its own internal real time clock using NTP. The eduWOSM web GUI and the WOSM keypad and the μManager interface update each other in real time, so that the eduWOSM user interface can be arbitrarily distant from the microscope. This can be helpful for example in containment facilities, for doing microscopy inside an environmental chamber, or for remote teaching. During the recent COVID pandemic, our undergraduate students were able to control their eduWOSMs from their homes, including internationally. There is nonetheless no requirement for an internet connection - imaging and data collection are fully functional with just a network cable between the PC and the controller. For both local and remote use, full digital control of every microscope parameter is implemented via the eduWOSM scripting language, a text-based command set that is optimised for speed (**Supp. Table 1**). The language has a rich set of intuitive commands that can be sent to the microcontroller from the web GUI, from a simple terminal application (like *PuTTy*), from μManager (via a second Telnet connection), or via the keypad. Commands can be combined into scripts. Loop timing has microsecond-level repeatability. Macros can trigger other macros and up to 8 macros can run simultaneously. Remote and local control update each other in real time and can be run simultaneously.

## Conclusions

The eduWOSM can bring research-quality optical microscopy into the classroom, with substantial benefits to students^18,19^, including the chance to learn more expert-like thinking^20^. In practice we are finding the eduWOSM useful not only for teaching (***Figs. 2-4***), but also for research. EduWOSMs are routinely used for research in our own labs, where their minimal footprint, user-friendly operation and high stability bring new possibilities - for example several eduWOSMs can be run in parallel, by a single operator, on a standard lab bench. We emphasise that the project is fully open source - software, gerber files for electronics, STL files for 3D printing and CAD source and engineering drawings for CNC are available from WOSM.net, which includes links to explanatory videos and exemplar movie data on the eduWOSM YouTube channel^21^.

## Materials and Methods

Many components for the eduWOSM use 3D printing in PLA plastic, including the keypad, electronics box, port covers on the microscope, and the sample hood. Three components, the main chassis, the sample mount bracket and the LED heat sink, require CNC machining in aluminium.

### Chassis

The eduWOSM chassis is CNC-machined aluminium (Width: 220 mm, depth: 170 mm, height: 100 mm). By default the chassis accepts an oil immersion, infinity corrected, 100x, 1.3NA Plan Fluor Nikon objective lens that screws directly into the chassis (Nikon thread 25mm × 0.75mm metric). The upper face of the block has threaded holes to mount the sample stage and bright-field illumination module and may be readily adapted for other user-defined hardware e.g. magnetic tweezers or micro-injection.

### Sample stage

A commercially available 3-axis stage (Thorlabs MT3) is used for sample XY and focus, providing 13 mm of sample movement in each axis. A stepper motor-based actuator (Newport TRA12PPD) is used to drive the Z-axis, allowing for fully automated 3D/4D imaging in Micromanager. In single-stepping mode this motor gives repeatable 97.7nm focus steps. Backlash compensation in firmware makes sample focus smooth, predictable and repeatable. Positional drift is so slight that single fluorophore tracking over tens of minutes can be done routinely without drift correction.

### Dichroic block

The dichroic mount holds an upper dichroic filter and a lower broadband mirror. The assembly is 3D printed in two parts, which are then screwed together. Both the upper mirror (Chroma or Semrock quad LED dichroic) and the lower mirror are held in place using adjustable 3D printed plastic springs and aligned by adjustment of the underlying 3-point mounts using grub screws. The sides of the block are intrinsically sprung so that the block fits snugly into one of the ports of the block and slides firmly into place, aligned to the optical axis, beneath the objective. Typically, alignment is done just once, at installation.

### Light engine

For fluorescence imaging, a custom 4-colour light engine provides excitation light in UV, blue, green and red (respectively 385-410 nm, 460-490 nm, 545-570 nm and 625-645 nm). The CNC-machined aluminium heat sink permits the LEDs to be run sustainedly at 0.7A without significant heating at the LED. Typical fluorescence microscopy applications require only a tiny fraction of full power. The light engine is small enough to fit inside the chassis. Three dichroic mirrors combine the output of the 4 LEDs. An f=19 mm achromat collector lens produces a 5x expanded LED image at the back focal plane of the objective lens. LED outputs are superimposed at assembly time and require no further alignment.

### Transilluminator

For brightfield imaging a small white LED is built into the sample hood (below) and illuminates a 3D-printed diffuser, which gives sample illumination up to 0.5 NA. The resulting lens-free, low NA condensor provides surprisingly good sample contrast. Transmission illumination does not need a bright source, so pulse-width control of a 3.3V logic line is more than enough to saturate the camera when recording the full field, unbinned, at 30 frames per sec. By altering the pulse width from 0 to 20 μs, the 50 kHz flashing of this LED provides sufficient power adjustment (equivalent to >10 bits of brightness adjustment). 50 kHz is fast enough to make pulsing/aliasing invisible until well below 1ms exposure time.

### Camera & adaptor

All our currently-recommended cameras (Teledyne/FLIR Blackfly S BFS-U3-50S5M-C) use the same Sony IMX264 monochrome sensor (2448 × 2048, 3.45um square pixels). The eduWOSM Nikon objective has a 100x magnification when the regular Nikon tube lens (f=200mm) is used. However here we use a tube lens where f=75mm. This brings two advantages over the regular tube lens. Firstly the optical pathway to the CMOS sensor is 12.5 cm shorter, keeping the instrument footprint small, and enhancing system stability. Secondly, and more importantly, the small pixels on the CMOS sensor are sampling the image at very close to the ideal sampling rate (92 nm pixels). The camera field is 225μm × 188μm. The real magnification is reduced to 37.5x, but the numerical aperture remains the same at 1.3, so there is no loss in actual optical resolution. Tests with sub-resolution tetraspeck beads have shown that little or no chromatic aberration results from this change in tube lens. Where a larger field is needed we use the 20x 0.9NA air objective lens (Nikon, order#: MRH00205), effectively 7.5x magnification. Here the full camera field sampled is 1.14 mm × 0.9 mm (0.46 μm pixels).

### Microcontroller

The microcontroller is a 252MHz PIC32MZ-EF core running custom firmware for microscope hardware control. A 100Mbit Ethernet module, with onboard W5300, controls TCP/IP comms as well as network data buffering and handshaking, independent of the main microcontroller. This module provides up to 8 sockets for Ethernet traffic which are used for the following network protocols: 2x HTTPD (for browser-based microscope control), 1x HTTP client (for firmware updates), 2x Telnet (one user terminal, one for the μManager driver). A single socket is shared amongst short-lived protocols including DHCP, DNS and NTP. The microcontroller and the ethernet module communicate via a fast 16bit parallel bus, so network latency remains low. Typically a telnet request for a variable (e.g. get the focus motor position) is completed within 100 μs, including the request and the reply network transit.

A web server hosted on the controller board serves HTML5 pages for microscope control from any browser on the same LAN. Usernames and hashed passwords can be configured. From the browser, pages are continuously polling for any hardware changes (by AJAX), so that the web pages are updated in real-time to reflect any changes at the microscope. (e.g. user changes made via the keypad).

The eduWOSM scripting language is optimised for speed and timing accuracy and allows the user to control the microcontroller via macros/scripts. Text is interpreted on-the-fly, average command time is about 5 μs. Simple loops and variables are supported. Macros are stored on the FAT32 formatted SD card in simple and legible text format. (Command set reference: https://wosm.net/mcu/commands.php). The controller configuration itself is saved as a macro (config.wml) that runs at boot-up. Users can download updates from wosm.net, or the microcontroller can be configured to automatically download updates. Updates, consisting of new firmware images and new templates for the html5 pages, are downloaded to the SD card. The user is given the choice to flash the new firmware on the next reboot. Alternatively, installation of updates can be triggered from the keypad. The update process takes about 30s.

The controller has 26x lines of TTL I/O available labelled A-Z. These are multi-function lines that can each be configured to be logical input or output. Two of the lines can be configured as 16bit counters.

5x PWM lines can be configured for r/c servo motor control (20Hz duty cycle) or LED brightness PWM control (50kHz duty cycle).

Up to 4 stepper motors can be controlled, with configurable stepping rate, accel/decel rates, and backlash compensation.

The microcontroller temperature bus can monitor up to 5x DS18B20 temperature probes).

A Raspberry Pi compatible connector allows for microscope control over SPI (e.g. from Python scripts on the Pi).

1x USB and 3x RS232 lines are present in hardware, but currently (Aug 2024) have no support in firmware.

### Powerboard

The powerboard provides power via an ATX connector, as found on most PC motherboards. The powerboard provides power for the microcontroller and LED driver modules and stepper motor drivers. An 8-channel i2c DAC provides 16bit control for LED modules. On the eduWOSM four LED driver modules (Recom RCD-24-0.70), provide 16 bit brightness/current control for the LEDs in the light engine (0 to 700mA). Each DAC channel has an associated TTL on/off control line for fast TTL-based on/off, via n-channel MOSFETs. The LED driver module has a turn-on delay of about 50 μs (the time from TTL rise to light output). Turn-off delay is about 200 μs.

### Keypad

The eduWOSM keypad has an LCD-based menu system for hardware control. The home screen has 8 user-configurable fields. Users can turn components on and off using hardware buttons and fine-adjust settings with single-digit precision using the rotary encoder. Settings are stored on powering-down components. The rotary encoder switches straightforwardly to control different microscope functions, for example sample focus, stepper motors, servos, adjusting DAC output etc. Sub-menus have more detailed hardware information, and functions such as web update and firmware flash. Five macro buttons are available to run user-defined macros “keypad1.wml” to “keypad5.wml”.

### Assembly and alignment

**Supp. Table 2** is a bill of materials for the eduWOSM, with prices correct in Aug 2024. Step-by-step assembly instructions are given in **Supp. File 2**. Please visit WOSM.net for further information and updates.

## Acknowledgements

The eduWOSM has been developed as a spin out from the microscope engineering element of a Wellcome Investigator Award to R.A.C. [220387/Z/20/Z].

