## Supplemental File 2 for "The eduWOSM: a benchtop advanced microscope for education and research"

N.J. Carter[+] D.S. Martin[*], J.E. Molloy[+o] & R.A. Cross[+]
[+] Centre for Mechanochemical Cell Biology, Warwick Medical School, Coventry CV4 7AL, UK
[o] Single Molecule Enzymology Laboratory, The Francis Crick Institute, London NW1 1AT, UK
[*] Department of Physics, Lawrence University, Appleton, United States

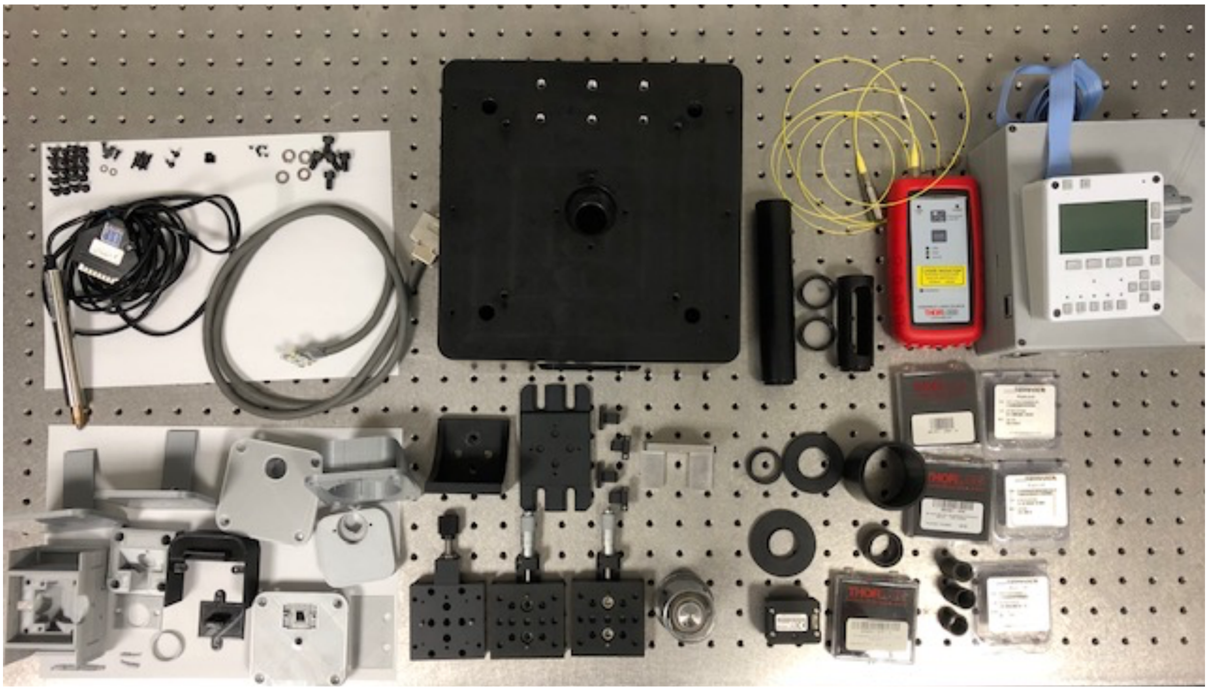

| **eduWOSM Parts List**<br>*please refer to Supp. Table 1 (BoM) for updated price guide* | | | |
|---|---|---|---|
| **Source** | **Part Number** | **Quantity** | **Total Price** |
| **Alignment** | | | **($1176.73)** |
| Thorlabs handheld laser | HLS635 | 1 | $680.65 |
| Thorlabs 2 m SM fibre | P1-630A-FC-1 | 1 | $70.07 |
| Thorlabs fibre collimator | F260FC-B | 1 | $160.15 |
| Thorlabs fibre adaptor | AD1109F | 1 | $32.74 |
| Thorlabs Nikon to lens tube adaptor | SM1A11 | 1 | $21.86 |
| Thorlabs alignment disk | DG10-1500-H1-MD | 2 | $69.82 |
| Thorlabs slotted lens tube | SM1L30C | 1 | $71.96 |
| Thorlabs lens tube | SM1E60 | 1 | $48.70 |
| Thorlab 0.5” to 1” lens tube adapter | SM1A6 | 1 | $20.78 |
| **eduWOSM** | | | **$5,448.43** |
| Thorlabs Nylon-tipped setscrews | SS3MN4 | 1 | $12.78 |
| Thorlabs 1” lens tube 0.5 inch long | SM1L05 | 1 | $12.97 |
| Thorlabs 1” lens tube 2 inches long | SM1L20 | 1 | $17.00 |
| Thorlabs 0.5” lens tube 0.5 inch long | SM05L05 | 1 | $14.52 |
| Thorlabs 0.5” lens tube 1 inch long | SM05L10 | 2 | $31.10 |
| Thorlabs 30 mm achromat, 0.5 inch dia | AC127-030-A | 1 | $55.19 |
| Thorlabs 75 mm achromat, 1 inch dia | AC254-075-A | 1 | $81.16 |
| Thorlabs 1” square dielectric mirror | BBSQ1-E02 | 1 | $77.35 |
| Thorlabs stage with lead screw adjuster | MT1B/M | 1 | $236.98 |
| Thorlabs stage with micrometer | MT1/M | 2 | $631.96 |
| Thorlabs XYZ base | MT401/M | 1 | $25.10 |
| Thorlabs Z mount | MT402 | 1 | $54.65 |
| Thorlabs side micrometer mount | MT405 | 2 | $122.84 |
| Newport motorized actuator | TRA12PPD | 1 | $788.00 |
| FLIR Chameleon3 2k x 1.5k USB 3 camera | CM3-U3-31S4M-CS | 1 | $489.00 |
| (FLIR Chameleon3 1.3k x 1k USB 3 (150 fps)) | (CM3-U3-13Y3M) | (1) | ($369.00) |
| (FLIR Blackfly S 2k x 1.5k USB 3 (118 FPS)) | (BFS-U3-32S4M-C) | (1) | ($765.00) |
| Nikon 100x 1.3 NA Plan Fluor objective | N100X-PFO | 1 | $2060.00 |
| Luxeonstar Saber Z4 with UV 395 nm, Blue, Lime, Red LEDs. | SZ-04-U6H3H9H6 | 1 | $117.00 |
| Semrock BrightLine filter set | LED-DA/FI/TR/Cy5-B-000 | 1 | $1468.20 |
| (Chroma multi LED filter set as alternative) | (89402-ET) | (1) | ($1200.00) |
| M3x6 mm set screws | | 4 | |
| M4x10 mm hex drive flat-head screws | | 20 | |
| M2.5x5 mm socket head screws | | 8 | |
| M6x6 mm socket head screws | | 5 | |
| M3x6 mm button head screws | | 4 | |
| M3 washers | | 2 | |
| M6 washers | | 4 | |
| WOSM Block | | | |
| WOSM Electronics | | | |

***Notes***

A. Wear gloves (nitrile or latex) when handling optics; only handle optics by edges, never by surfaces.

B. Suggested alignment laser: Thorlabs HLS635 handheld laser source + P1-630A-FC-1 1m single mode fiber + F260FC-B fiber collimator + AD1109F collimator adaptor. This alignment laser will have a 3 mm dia red beam, and be easily moved from alignment port to alignment port.

C. Basic alignment laser safety:
   a) The laser is roughly as bright as the sun, midday in summer.
   b) Do not shine beam into someone else's eye. Even by accident.
   c) Do not stare into beam. If beam enters the eye, blink and turn head. The beam will dazzle, but should not cause permanent damage any more than the sun does.
   d) Do not wear reflective jewellery on hands or fingers while aligning beam; reflections from mirror-like surfaces are just as dangerous as the direct beam.
   e) Turn alignment laser off between uses.

1) **Order the chassis** from a machining contractor. Get it anodized.

2) **Construct the dichroic.**
   a) **Build the alignment tool:** SM1A11 adaptor, DG10-1500-H1-MD alignment disk, SM1L30C

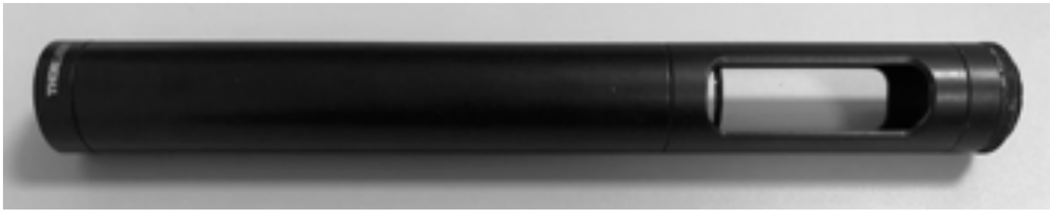

   slotted lens tube, SM1E60 lens tube, DG10-1500-H1-MD alignment disk.
   b) **Print the dichroic parts**: 1in_mirror_mount, ccd_mirror_spring, dichroic_mirror_spring, dichroic_mount, emission_filt_cover, filter_spacer, imaging_alignment_port, mirror_mount_base, LED_path_support from the LED illuminator, and the 2in_support_camera from the camera mount.. 0.2 mm layers through a 0.4 mm nozzle on a Ultimaker 3 has worked for us; ~ 24 hours for all part to print.
   c) **Place the imaging mirror**
      i) Screw M3 nylon-tipped screws into 1 inch mirror mount until just below surface.

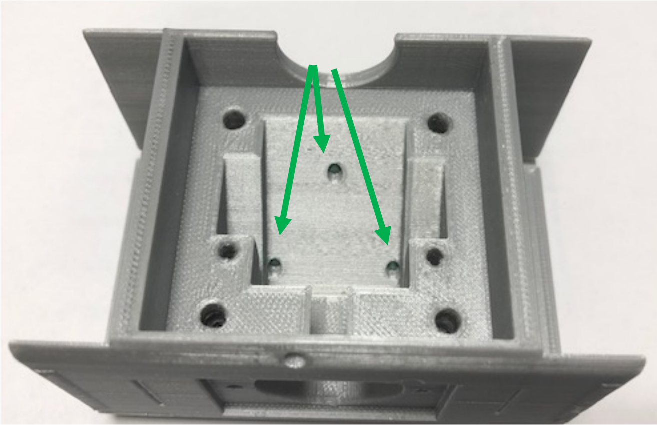

      ii) Insert the ccd mirror springs into the 1 inch mirror mount.
      iii) Slide 1 inch mirror (BBSQ1-E02) into 1 inch mirror mount, behind springs. Mirror surface should face out.

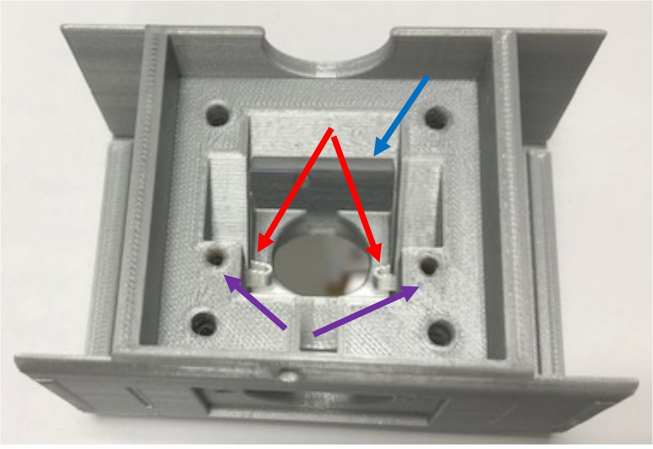

      iv) Use M3x6mm set screws to tighten springs onto mirror. Mirror should be held in place, but springs should not flatten onto mirror.
      v) Stack the dichroic mount onto the 1 inch mirror mount, oriented so that LED light-path channels line up.
      vi) Use M4 flat-head screws to bolt dichroic mount onto 1 inch mirror mount.

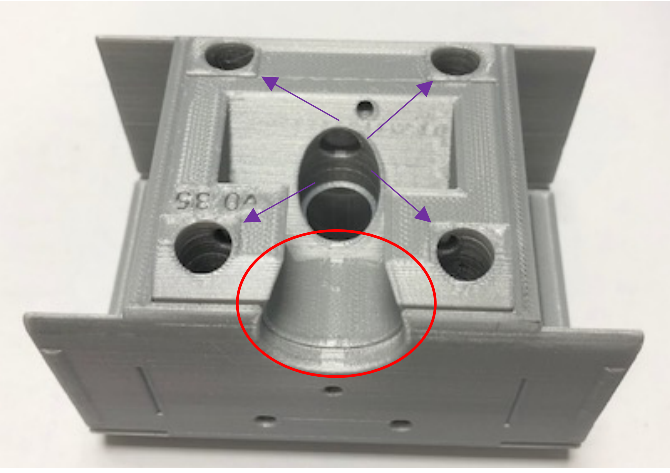

vii) Insert dichroic stack into block, looking from end to make sure camera port is centred.

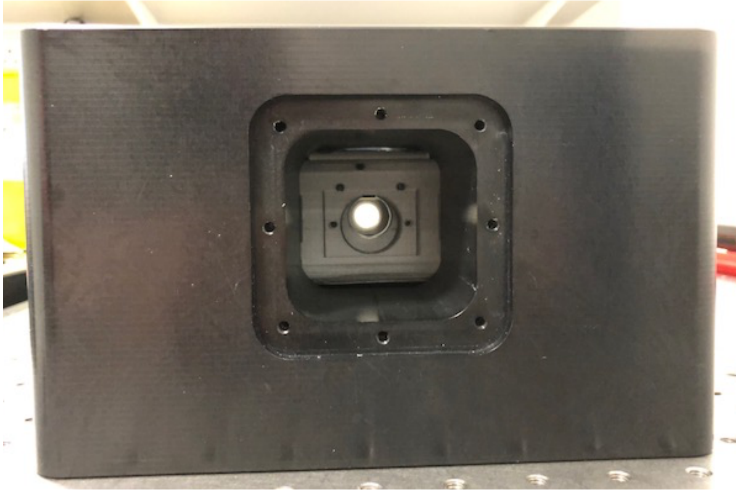

### d) Align the imaging mirror

i) First, build camera mount as follows:

(1) Screw together tube lens housing: SM1L05 into SM1L20, with exposed threads on the SM1L20 tube.

(2) Slide tube lens housing into the 1 inch camera support 3-D part SM1L05 tube is flush with the plastic. NOTE: This is backwards relative to the final camera construction; it is for alignment only.

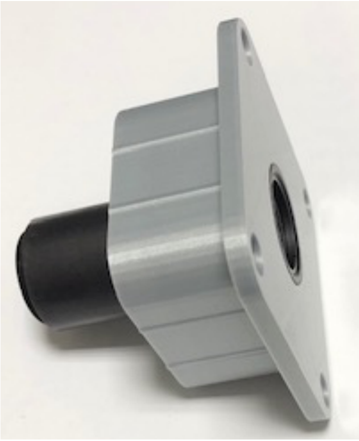

(3) Slide camera mount into the block and secure using M4x10 mm flat head screws.

ii) Attach SM1A6 adapter to end of camera mount and attach alignment laser to SM1A6 adapter (note: screw on AD1109F adapter, then attach fibre, to avoid unnecessarily twisting the fiber). Tighten SM1A6 against a SM1 retaining ring. Fibre laser should now be directed along the imaging path, from the camera back to the dichroic mount, and then back to the sample. Do not turn laser on yet (it will shine directly up, toward eyes).

iii) Screw alignment tool into objective port on the WOSM block.

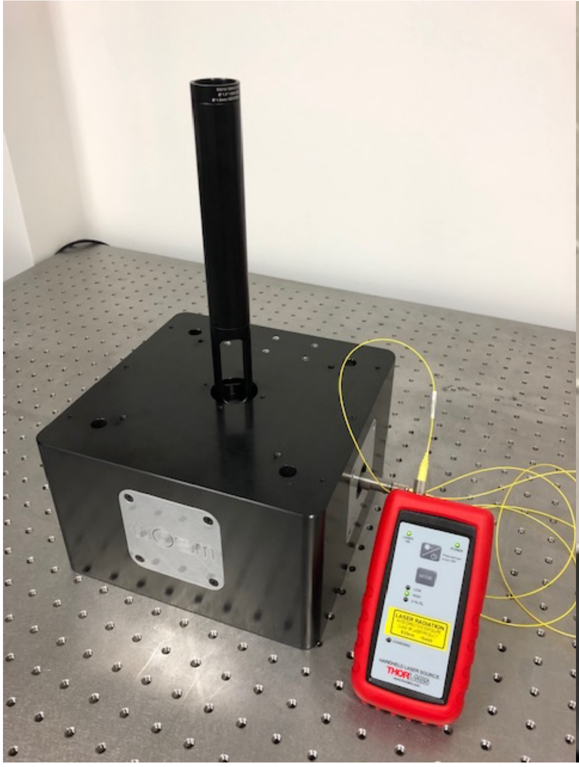

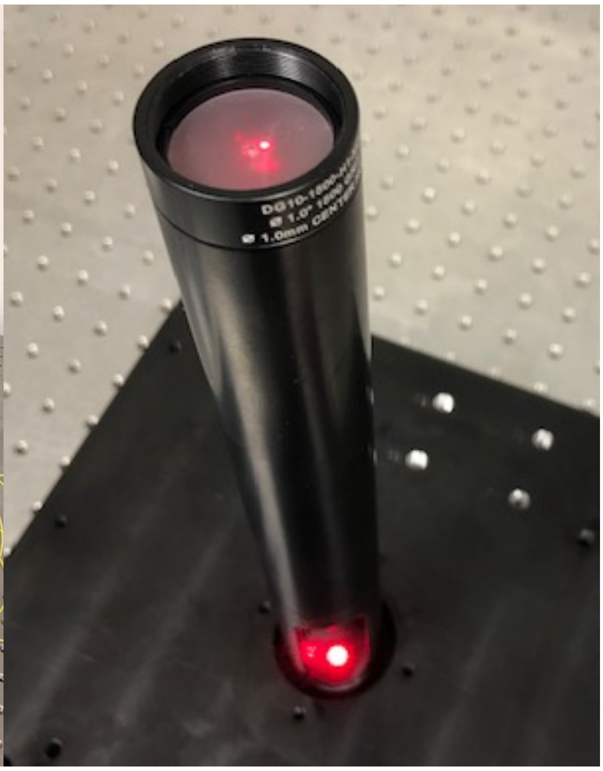

iv) Turn on alignment laser (lowest power); beam should be visible on lower ground glass disk of alignment tool.
   (1) If beam is not visible, turn laser off and check for obstructions. Make sure dichroic is oriented so that the 1 inch mirror faces the camera port.

v) Align the mirror:
   (1) Using a 1.5 mm hex key, turn each of the M3 nylon-tipped screws CW, in turn, until beam just moves on ground glass plate. If the beam moves immediately, turn CCW until beam stops moving; this indicates the screw is not touching the mirror. GOAL: have 1 inch mirror resting on the three M3 nylon-tipped screws rather than on plastic.
   (2) Mirror is on a tripod of screws (U/D, L/R); adjust correct screws until beam is centred on the hole in the bottom ground glass disk.

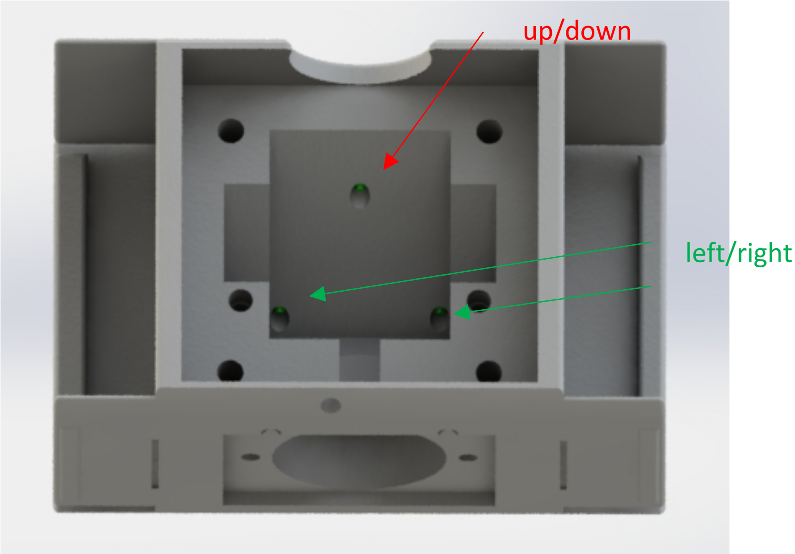


(3) Adjust screws until beam is centred on the hole in the upper ground glass disk of the alignment tool. The beam may not be centred on the bottom hole. DO NOT EYEBALL BEAM COMING OUT OF UPPER GLASS DISK.

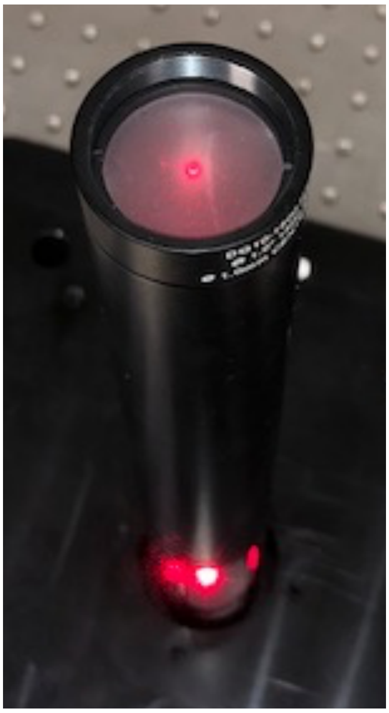

(4) IF beam is off to the left or right, the laser is not aligned – check camera mount to make sure every part is tightly attached.

(5) If beam is too high on the lower ground glass disk, turn each M3 screw the same amount CW to move entire mirror forward. 1/8th of a turn at a time works pretty well.

(6) If beam is too low on the lower ground glass disk, turn each M3 screw the same amount CCW to move entire mirror backward. If the mirror cannot be lowered enough, the 1 inch mirror mount will have to be reprinted.

(7) Repeat steps 3-6 until beam goes through the centre of the lower and upper disks in the alignment tool. The 1 inch mirror is now aligned.

vi) Remove dichroic block from the WOSM block, and remove the dichroic mount by loosening the M4 screws.

**e) Place the dichroic mirror:**

i) Screw M3 nylon tipped screws into dichroic mount until just below surface.

ii) Insert dichroic mirror springs into dichroic mirror.

iii) Slide dichroic mirror, SEMROCK LED-DA/FI/TR/Cy5-B, behind springs. NOTE: Reflective surface is indicated by a small engraving on the dichroic mirror. Reflective surface should face up.

iv) Use M3x6 mm set screws to tighten springs onto dichroic mirror. Mirror should be held in place, but springs should not flatten onto mirror.

v) Rebuild dichroic block with the 1 inch mirror mount, and insert dichroic block into WOSM block, oriented correctly.

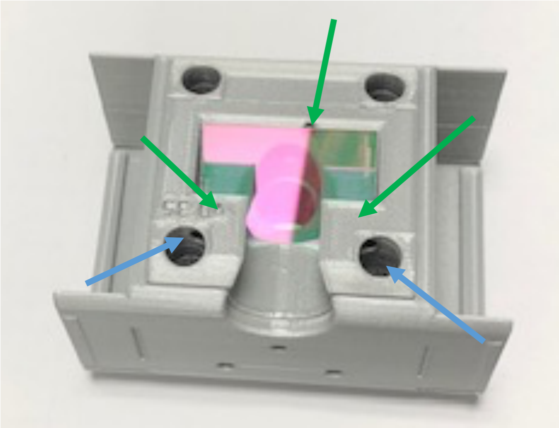

**f) Align the dichroic mirror:**

i) Insert a 2.5-inch long ½ inch lens tube through the illumination alignment port, (use the SM05L10 x2 and SM05L05 parts), with threads facing in.

ii) Screw alignment laser into ½ inch lens tube.

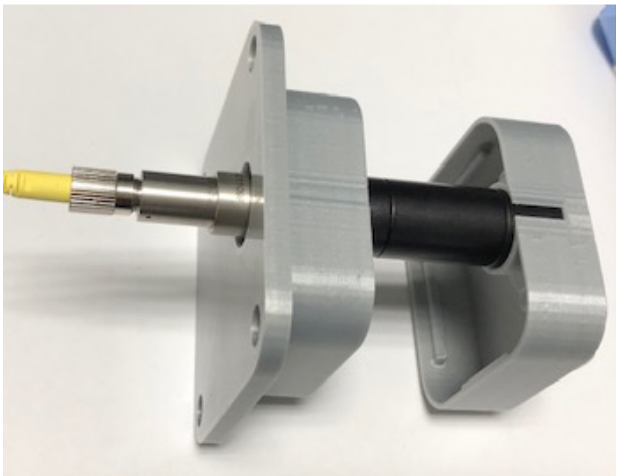

iii) Insert LED path support into illumination port of the WOSM block, and then insert illumination alignment port. The ½ inch lens tube should slide into LED path support and be pushed all the way until it hits the dichroic block. Screw illumination alignment port into block using M4x10 mm hex drive flat screws and turn on laser.

iv) As in *dv1-6*, turn M3 nylon tipped screws in dichroic mount to centre alignment laser on bottom and top holes of the alignment tool.

g) **Return alignment laser to camera port.** Remove dichroic mirror alignment port, and revisit *dv3-6* to make sure imaging mirror is aligned through the dichroic.

h) **Insert Semrock emission filter** into filter slot in dichroic mount (using arrow to align correctly), add filter spacer, and secure using emission filter cover and M3x10 mm flat head screws.

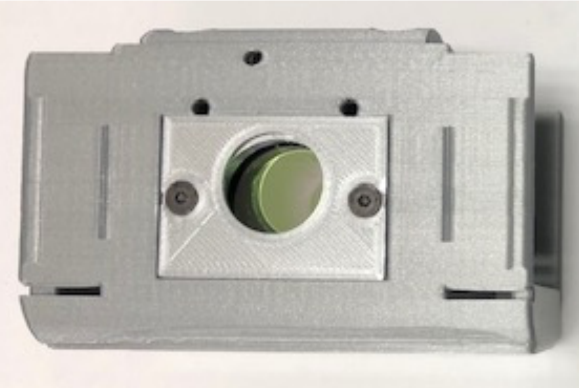

i) **Dichroic construction is complete!** For future use, centring can be done by eye using lines on dichroic block or using end-stops on the side caps.

### 3) Construct the LED illuminator:

a) Print remaining LED illuminator parts: Z4 and tube mount v2, cableclip.
b) Solder wiring for power onto the Saber Z4 chip as described in the Z4 documentation.
c) Put Z4 chip into the Z4 and tube mount v2 part facing in, and secure using M3x5 mm button head screws. Use cableclip and M6x6 mm screw for cable strain relief.

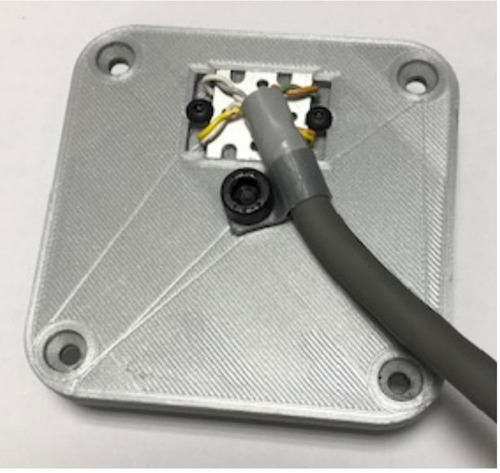

d) Insert 30 mm f achromat lens AC127-030-A into 1 inch long ½ inch diameter lens tube SM05L10 with rounder side facing away from external threads and secure using retaining ring.
e) Screw lens-containing lens tube into a second SM05L10 lens tube and SM05L05 lens tube as shown below.
f) Push lens tube into Z4 and tube mount part, ensuring that the 0.5 inch lens tube is closest to Z4.

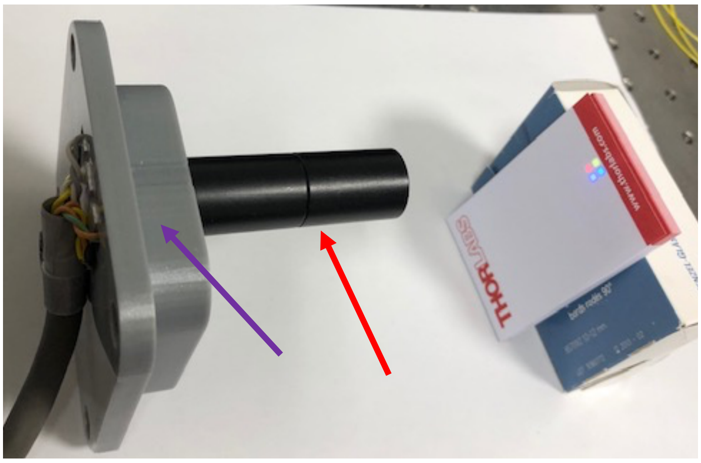


g) Put the SEMROCK excitation filter into the LED path support (oriented as shown on SEMROCK part) and secure using M3x5 mm button head screws and washer.

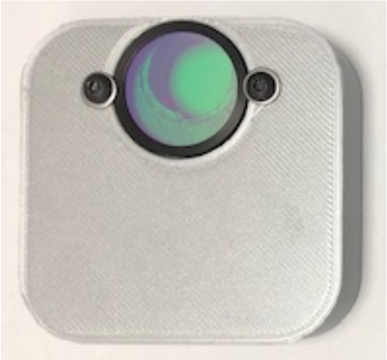

h) Slide LED path support with filter facing the dichroic into the illumination port of the WOSM block, and then slide the Z4 and tube mount in, lens tube forward.
i) Test alignment by turning on all four LEDs using WOSM. Four square LED images should appear, centred and in focus, on the lower ground glass disk of the alignment tool. If not centred, check Z4 placement. If not in focus, check 30 mm focal length lens placement.

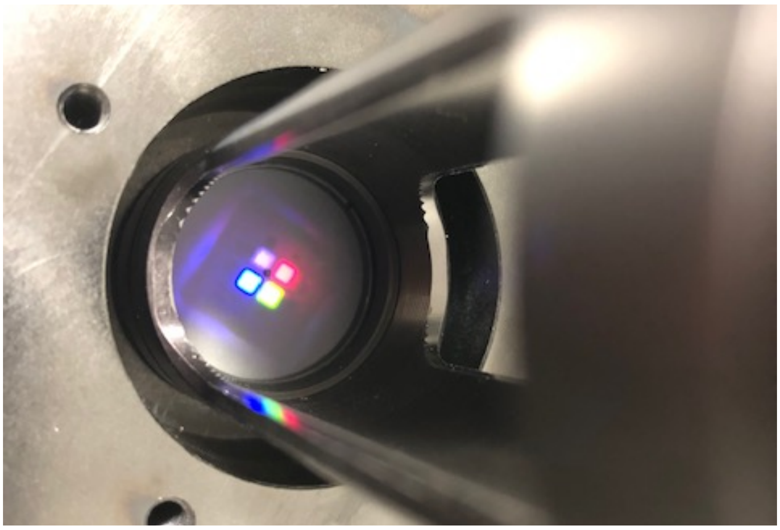

j) LED illuminator construction is complete.
k) ALTERNATIVE: Build LED light engine using Knight Optical dichroics 420FDL (10x12 mm), 565FDL (14x12 mm) and 635 FDL (10x12 mm) [potential alternatives: Comar Optics 454 IY, 560 IY, 650 IY]. Use Thorlabs AC127-019-A 19 mm lens as a condenser.

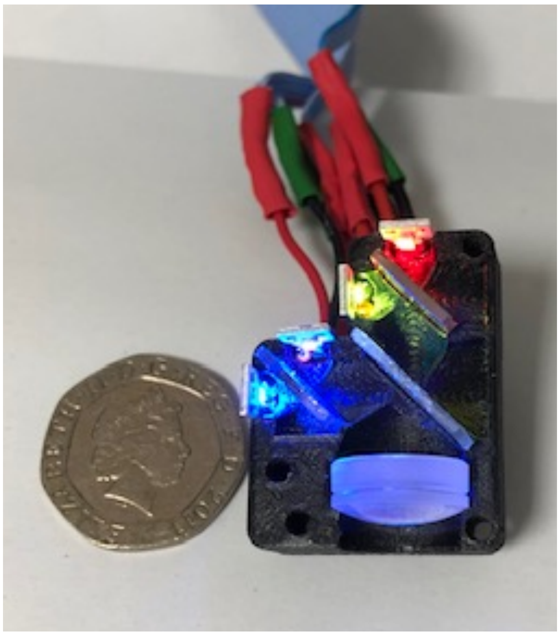

4) **Construct the camera mount:**
   a) Remove Thorlabs 1” lens tube assembly from camera mount and remove SM1A6 adaptor.
   b) Place 75 mm F achromat into the 0.5 inch lens tube, round-side out, and secure with retaining ring. NOTE: this leaves the back of the lens 71.3 mm from the camera chip of the Chameleon3. The back focal length is 70.3 mm, close to the actual distance.
   c) Put lens tube assembly back into 1 inch camera mount, lens inward, with 2 inch tube flush with plastic.
   d) Screw c-mount to SM1 lens tube adaptor SM1A9 onto end of camera mount.
   e) Screw camera onto adaptor and re-insert into WOSM block.

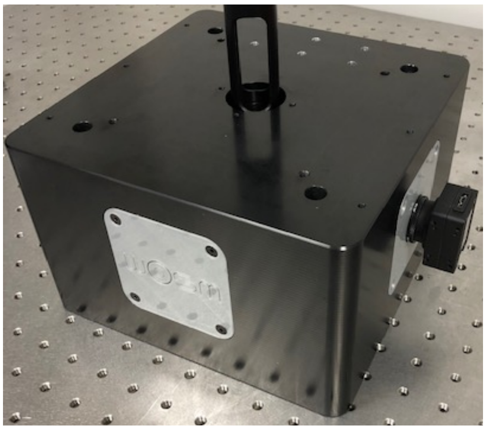

   f) Camera mount construction is complete.

5) **Construct the sample stage:**
   a) Print the MT1_Z_shim, light_blockv2, light_curtainv2.
   b) Mill the MT3_sample_bracket.
   c) Remove micrometer from Z-axis.
   d) Move Z-axis micrometer attachment from top to side using MT405 PHOTO.
   e) Attach motorized actuator to Z-axis.

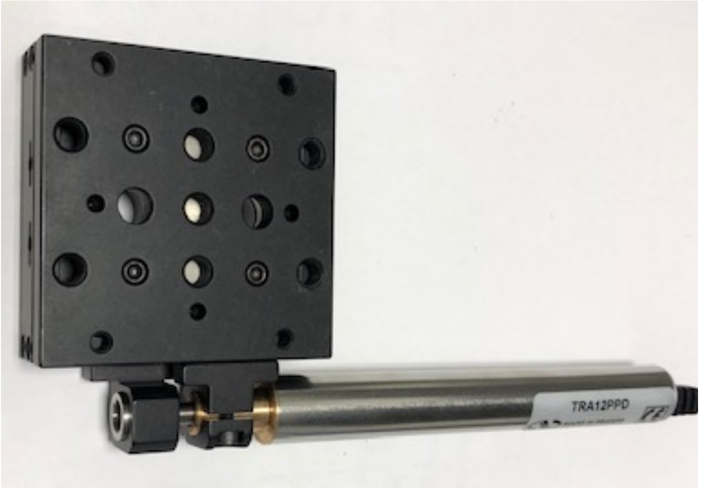


   f) Attach to the angle plate (MT405) via the shim, one set of holes lower. Be sure to bolt onto the side of the stage that stays fixed when the actuator is moved.
   g) Move Y-axis micrometer and attachment from end to side using MT405.
   h) Attach the MT3 sample bracket to Z-axis of stage using two M6x6 mm socket head screws.

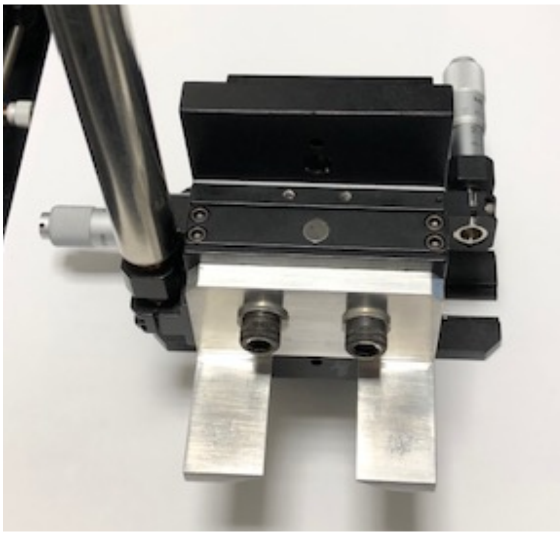

i) Remove alignment tool.
j) Screw objective into place.
k) Attach sample stage to WOSM block using four M6x6 mm socket head screws.

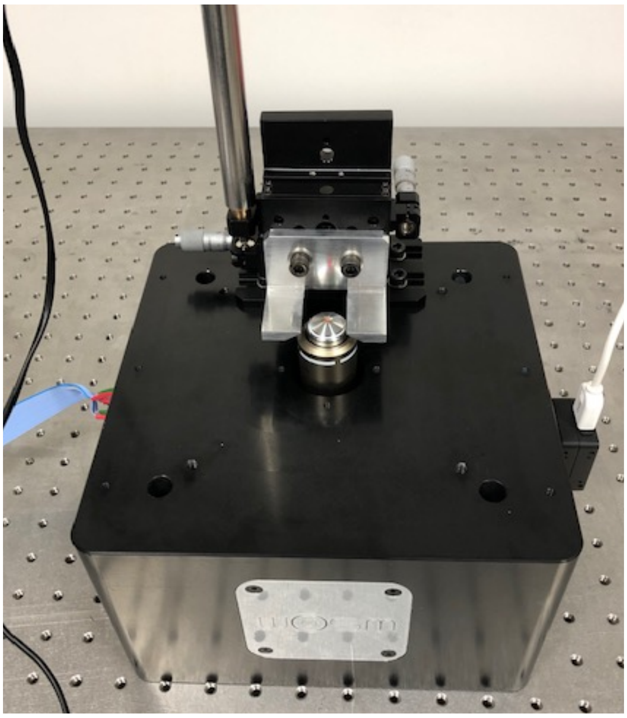

l) Screw M2.5x5 mm screws into light block and light curtain.

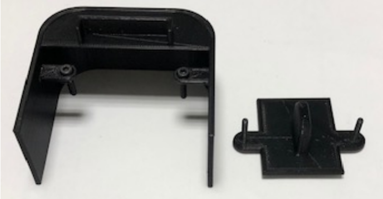

m) Stage construction is complete.

6) **eduWOSM construction is now complete.** Plug everything into WOSM electronics box and computer, try samples using micro-manager. Suggested test sample: Molecular Probes TetraSpeck Fluorescent Microspheres Size Kit, ThermoFisher T14792.
