## Supplemental Table 1 for "The eduWOSM: a benchtop advanced microscope for education and research"

**System**

| Command | Arguments | t (µs) | Description |
|---|---|---|---|
| sys_reset | | | System reset |
| sys_poweroff | | | Will power off the ATX power supply. NB. Jumper J? can override this. |
| sys_interf | int | (4) | Read/write the system interface bitfields:<br>bit0: ( 1) ethernet<br>bit1: ( 2) usb (not supported yet...)<br>bit2: ( 4) keypad: lcd, buttons and leds<br>bit3: ( 8) rotary encoder (usually on keypad)<br>bit4: ( 16) PSU ATX power control<br>bit5: ( 32) PSU board analogue out<br>bit6: ( 64) DS18B temperature modules<br>bit7: (128) Addressable LED bus |

**General**

| Command | Arguments | t (µs) | Description |
|---|---|---|---|
| delta | [all]<br>[clear] | | Delta is a "what's changed" call. Used by an interface, controller software or a web page (eg. a request for "/delta.json") to get updates on any hardware changes. A reply to a delta command will include the command name (command details throughout this page) followed by the current parameter value. A single parameter change is reported for each delta call.<br>eg.<br>delta<br>dig_in 0x00000009<br>delta<br>sys_unixtime 1567676848<br>If there's nothing to report, a blank line is returned. The controller maintains an internal list of what has been reported to each interface (http, usb, driver telnet, user telnet, SPI A). This way a parameter update will only be sent once, to each interface.<br>The "**all**" argument will request to re-send all parameters (now in the hundreds) on subsequent calls to delta.<br>The "**clear**" will clear all the delta pending flags, so only newly changed parameters will be sent. |

**Timing**

| Command | Arguments | t (µs) | Description |
|---|---|---|---|
| sys_time | | 14 | Returns the system time (UTC). Read only. |
| sys_date | | 12 | System date. Read only. |
| sys_usec | | 2.5 | Returns the time since boot in microseconds. Useful for accurate timing in macros. |
| sys_usec_wait | time_usec | | [Macro-only] Pauses the macro until the system microsecond value (sys_usec) reaches time_usec. |
| sys_uptime | | 4 | Read only. How long since switch on. Days, hours, mins & secs. |

| | | | |
|---|---|---|---|
| sys_ntp_last | | | GMT string of most recent NTP check |
| sys_ntp_next | | | GMT string of the next NTP check |
| sys_ntp_now | | | Force immediate NTPD check. |
| sys_xtal_ppm | | | Crystal error "parts per million" relative to the NTP server references. |
| sys_unixtime | unixtime | 3 | Get/set the wosm unixtime. Returns the current UTC (seconds since start of 1970) |

**Temperature modules (DS18)**

The number X in the command refers the module index (0 to number of modules - 1).

| Command | Arguments | t (µs) | Description |
|---|---|---|---|
| temp_ser | X [serial] | 8 | Get/Set the serial (hexadecimal) number for this module. |
| temp_ref | X [setname] | 2.5 | Get/set the friendly name (reference) for this module. Up to 7 characters.<br>eg. temp_ref 4 room |
| temp_off | X [offset] | 6 | Get/set the temperature offset. For temperature offset calibration<br>eg. for a +1°C offset correction "temp_off 0 256". Offset values are saved in the module's limited storage space, so it should be retained if you move modules later. |
| temp_val | X | 2.7 | Get the last temperature reading from module. Signed integer value returned with 256th degree units (ie. divide by 256 to get degrees. |
| temp_d5m | X | 2.8 | Get the temperature change over the last 5 minutes. (256th degree units). |
| temp_deg | X | 6.5 | Get the last temperature reading from module. Value returned in degrees with single decimal place (eg. "23.42"). |
| temp_col | X [colour] | | GUI colour used for this temperature module. In hex<br>eg. temp_col 0 2020D0<br>Would be parsed as "#2020D0" in the html. For help choosing codes go to<br>http://html-color-codes.info |
| temp_set_all | signed float | | Correct the offset calibration for all modules to give the temperature you set here.<br>A useful set-up calibration. Have all modules running together for several minutes in the same temperature controlled environment, ideally next to a trusted thermocouple. Or what about in a bag in melting ice? |

| | | | |
|---|---|---|---|
| | | | eg. "temp_set_all 21.3" will correct all module offsets so they all read out 21.3°C. |
| temp_log_clr | | | Zeros the temperature log buffers. |

**Addressable Serial LEDs**

Commands to control a serial run of addressable LEDs.

| Command | Arguments | t (µs) | Description |
|---|---|---|---|
| led_conf | n=numLEDs<br>T0H=n1<br>T0L=n2<br>T1H=n3<br>T1L=n4 | | Set the number of LEDs one the line with "n=??".<br>Bit timing can be adjusted by setting TL (time low) and TH (time high) to read ones and zeros on the bus. You can get these from the datasheet of your RGB or RGBW led. Default timings work for WS2812B LEDs, one of the more popular types. The timing numeric defines tenths of a microsecond. eg.<br>led_config ch=0 n=25 T0H=3 T0L=8 T1H=6 T1L=5<br># write to 25 LEDs<br># zero timing (0.3us high, 0.8us low)<br># one timing (0.6us high, 0.5us low) |
| led_buff | RGBval<br>[buff write mask] | | Setup the buffered values to send to the LEDs. This writes RGB(W) values to the buffer, **not** to the LEDs. This is where you define the colours and relative brightnesses of the LEDs (for overall brightness use the led_bright command).<br>The **buff write mask** sets the LEDs to write the new value (bit 0 = first LED, bit 1 = second ... etc).<br>eg.<br>led_buff 0xFF00FF 0x000000010 # make LED number 9 purple |
| led_bright | 0 - 255 | | This is the brightness multiplier (0 - 255) is applied to the led buffer before sending (dimmest 0 to 255 brightest). |
| led_mask | [ masknum / + ] | | Select which mask to use, by number (0 to 5) or select the next mask with "+".<br>led_mask 3 |
| led_mask_def | [n=masknum]<br>[ref="nameit"]<br>[mask=newmask] | | Set/get the LED mask. Define any one of 5 masks (n=1 to 5). Masks define which LEDs are active (bit 0 = first LED, bit 1 = second ... etc). The ref is just a friendly name for the mask, shown in the UI and keypad, 7 characters max. Mask n=0 is always "All LEDs on" ie. 0xFFFFFFFFFFFFFFFF. eg.<br>led_mask_def n=1 ref="BrightF" mask=0x0000FF00<br>#Just use LEDs 8 to 16<br>led_mask 1 #make mask 1 active |
| led_on | 0 or 1 | | LED array off (0) or on (1). |

**Digital lines in / out (TTL) eg. shutters, triggers**

The 26 digital lines are labelled A to Z.

| Command | Arguments | t (µs) | Description |
|---|---|---|---|
| dig_in | [a-z] | 2.1 | Read only. If specified, read from a single TTL in line. Returns 0 or 1. A return value of -1 means this line is not currently configured to be a ttl input. If no line is specified, a hex value representing all input lines is returned (bit 0 for line 'a', bit 1 for line 'b' etc.) |
| dig_wait | a-z<br>1 / 0<br>[t=??s] | | Read only. Macro only. Halt macro progress until the line reads high (1) or low (0). This will work for "TTL in" lines and "TTL out" lines.<br>Timeout: (v0.816 and later) If not specified, this command times-out after 1s. The timeout value can be entered in the command. eg:<br>dig_wait n 1 t=20s #wait 20s for line N to go high<br>dig_wait s 0 t=100us #wait 100us for line S to go low<br>By default, when this command times-out it will stop the macro. (see ... to prevent this...) |
| dig_out | [a-z] [0/1/2]<br><br>or<br><br>[val32 mask32] | 2.5<br><br>or<br><br>11.6 | Set a single output line low (0), high (1) or toggle(2) the output. eg:<br>dig_out c 1 #will set ttl output high on line C.<br>dig_out t 0 #switch off line t.<br>dig_out w 2 #toggle line w.<br>If no parameters specified a single hex value is returned, representing the out value of all digital output lines. |
| dig_mode | a-z<br>[setmode] | 4 | Set/get the line mode.<br>eg. "dig_mode a 4" will configure line A to be a "TTL out" line<br>(other modes available on certain lines, see web GUI) |
| dig_ref | a-z<br>[newname] | 3 / 5 | Get/set the reference/name for this digital line. |
| dig_hilo<br>dig_lohi | a-z<br>time<br>[nowait] | 5.5 | Accurately timed signal output commands. Immediately starts a pulse output on the line specified, enter the delay in the wosm time format. Either high(3.3V) to low (0V), or low to high. Useful for opening a shutter or switching on a laser for a accurately defined duration. eg:<br>dig_hilo q 1s #(open the shutter on line Q for 1s)<br>dig_hilo m 2us #(2 microsecond pulse on line m)<br>This should have microsecond repeatability. Try it on an oscilloscope and let me know....<br>Minimum pulse duration is 1µs.<br>If you add the "nowait" option, the command is non-blocking. After the first signal change, the macro continues with the next line. eg. in this command sequence<br>dig_hilo a 20ms nowait<br>dig_hilo b 20ms<br>The two output pulses are almost simultaneous, save for the approx 10us of interpreter time at the start of the signal on line b. |
| dig_all_modes | | 6.7 | Read only. List the modes of all lines A to Z. |
| dig_all_refs | | 6 | Read only. Get all reference names in one call. |

**MZ boards: line reference**

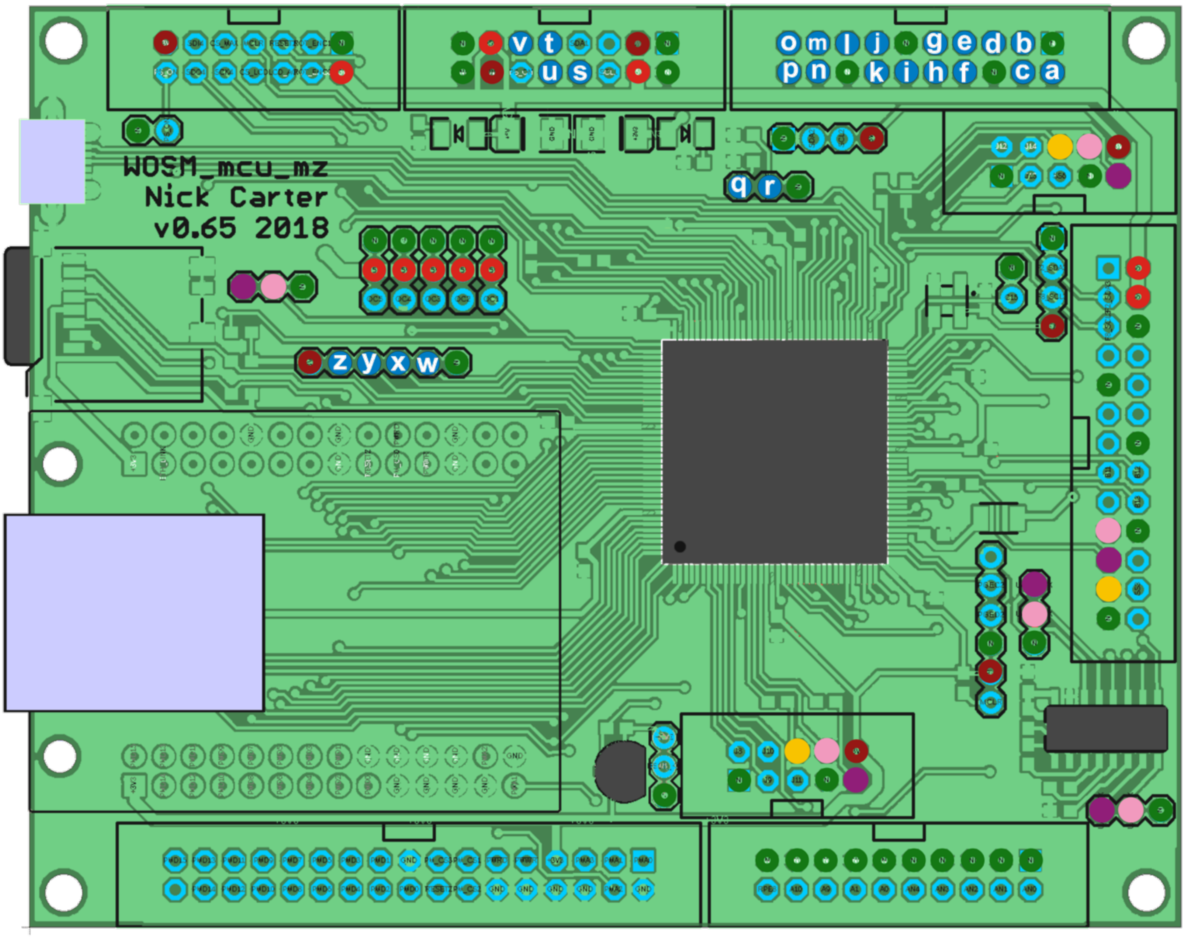


Key:

- Digital i/o
- GND
- +3.3V
- +5V
- Clock line (CLK)
- Serial out (SDO/TX)
- Serial in (SDI/RX)

**Pulse-width modulation: control R/C servos and LEDs**

By varying the on-time of rapidly repeating (50Hz to multi-kHz) digital signal can provide accurate analog-like control.
R/C servos use pulse width to control motor position.
LED brightness can be controlled by altering the duty-cycle of the pwm line. All commands must specify the channel argument ie. "1" to "5".

| Command | Arguments | t (µs) | Description |
|---|---|---|---|
| pwm_conf | ch<br>[mode=0/1/2]<br>[ref=aName]<br>[min=count]<br>[max=count]<br>[timeout=timeval] | | Configure the PWM mode for channels PWM1 to PWM5.<br>Modes: 0=line not used, 1= r/c/ servo, 2= LED pwm.<br>pwm_conf 2 mode=1 ref=Iris2 timeout=3s # r/c servo on channel 2<br>pwm_conf 4 mode=2 ref=Blue # Blue LED on PWM channel 4<br>Two modes are currently available:<br>**Mode 1**, typically for driving PWM based **R/C servos**. This will output a 50Hz repeating PWM signal with a default min equivalent to 1ms, and a max of 2ms. These min/max values can be overridden to give a servo more/less movement range. Take care not to strain or grind the gears, you can mangle them by going too far. **Mode 2**, typically for driving LEDs. This will output a 100kHz repeating PWM signal with a default min=0 and a max of 10µs. These min/max values can also be overridden. You can drive an LED directly from the line, but ensure an in-series resistor keeps the maximum current below 10mA (at 3.3V that would be 330ohms to be safe). You can drive higher currents by using the neighbouring 5Vpin, an in-series resistor and an n-channel MOSFET. |
| pwm_set | ch<br>[on=0/1/t/timeval]<br>[val=PWMval]<br>[percent=set_percent] | | Get/Set the pwm value. The pwm timing can be set as an actuatl timer count (val) or as a percentage.<br>In percentage terms "min" is 0%, and max is 100% (see pwm_conf for how to define min/max).<br>pwm_set 2 # get the on/off state and timer value of PWM2<br>pwm_set 3 on=4ms val=208 # PWM3 turn on for 4ms, set count to 208<br>pwm_set 5 on=t # toggle on/off PWM5<br>pwm_set 2 percent=50 # PWM2 duty cycle set to 50% |
| pwm_on | [bitmask] | | Single command to turn on/off all 5 channels by bitmask.<br>For channels 1,2,3,4,5 add 1,2,4,8,16 respectively.<br>pwm_on 0 # All pwm channels (1 to 5) OFF<br>pwm_on 31 # All pwm channels (1 to 5) ON<br>pwm_on 9 # PWM1 (+1) on, PWM4(+8) on, others off |

**Counters**

16bit counters will sum the signal pulses (3.3V TTL) on the line. Currently 16bit only. Should be capable of a few MHz (limits not tested yet..).
Counters are available on lines "q" and "r".
NB. You need to configure a line as a counter for these to work!

| Command | Arguments | t (µs) | Description |
|---|---|---|---|
| cnt_en | counter [0/1] | 2.6 | Counter enable. 0=disable, 1=enabled. The counter total is not changed so this command functions as a pause/continue.<br>eg. cnt_en m 1 #enable the counter on line m |
| cnt_en_all | 0/1 | 2.6 | All counters simultaneously. 0=disable, 1=enabled. |
| cnt_clr | counter | 2.4 | Zero the counter. |
| cnt_val | counter | 2.4 | Read only. Read the counter value. |

**Keypad, rotary encoder and LCD**

| Command | Arguments | t (µs) | Description |
|---|---|---|---|
| kyp_val | (read only) | 5.5 | Returns a 32bit hex value representing the keypad buttons and LED states. |
| kyp_rot | | 2.1 | Return the rotary encoder cumulative counter (signed 32bit int) |
| kyp_rot_expo | [rate] | | Zero means linear output from the rotary encoder. If non-zero, faster turns will take you much further. An acceleration effect, while maintaining good control when moved slowly. Test with a value of 10 and adjust up and down to your liking. The actual value varies depending on ticks/turn from your encoder and user. (Actual function is quadratic, exponential was too harsh). |
| kyp_led | LEDname [on/off/0-16] | 9.7 | Set the keypad indicator LED brightness (current) values (0=off, 1=dim, 16=bright). Slightly quicker for the "on" / "off" commands as there is no current set operation. The "on" command remembers the last brightness setting (defaults to 12 on boot.....check this...?).<br>Valid names for the LEDname are:<br>**usr1** spare led 1<br>**usr2** spare led 2<br>**m1...m5** macro leds, above the macro keypad buttons<br>**lcd** all lcd backlight leds<br>**lcdr lcdg lcdb** individual rgb leds if you have the RGB backlight |
| kyp_lcd_type | 0, 1 or 2 | | Which LCD screen hardware used in the keypad.<br>0 = No LCD used<br>1 = SP5 GFX1 LCD (older style keypads with vertically arranged macro buttons)<br>2 = EA-DOGL128 (default) |
| kyp_lcd_cont | 0 to 63 | | LCD contrast. Start with 30. Adjust to taste. |
| kyp_lcd_screen | [t=timeout] [small] [nowait] "text" | | Show a temporary LCD screen, with a specified timeout. Text is escaped before showing on the LCD. For a new line, use "\n". If you use small text, you can fit 8 lines of text, otherwise 4. |

| | | | |
|---|---|---|---|
| | | | eg. kyp_lcd_screen t=1s small "Alert!!!\nRead this quickly\nIt's only here for\n1 second."<br>If t is not specified, the default timeout is 2s.<br>**title** plots the first line inverted, as a heading, or a title.<br>**nowait** Macro progress will continue while the text is displayed. Without this the macro will pause until the screen times out. |
| kyp_lcd_text | [small] [inv] [upd] [x=0 to 127] [y=0 to 63] "escaped text" | | Show text on the current screen.<br>**small** for smaller text<br>**inv** to plot the text light and background dark<br>**upd** force immediate screen update (useful in fast mode, where the macro hogs the mcu so much that the screen does not get updated)<br><br>eg. kyp_lcd_text small x=20 y=33 "Yerluvinunclebert" |
| kyp_lcd_mode | mode number | | Get/set the LCD screen mode. |

**Stepper motors**

Motors are referenced in two characters referring to the host board (usually "m" for microcontroller board) and the motor number. The main mcu board can be configured with up to 5 servo motors (m1 to m5) and 4 sterpper motors (m6 to m9). Stepper positions and limits are defined in step units (steps or microsteps) and servos in pulse width units (in microseconds).

| Command | Arguments | t (µs) | Description |
|---|---|---|---|
| mot_ref | m1-4 [newname] | | Get / set the friendly name for this motor.<br>eg. mot_ref m4 Zdrive/td> |
| mot_stat | m1-4 | | Motor state, options are:<br>0 = Not available in this hardware config (dig io lines may need setting?) (default).<br>1 = Disabled / not configured<br>2 = Motor off. Stays off, does not respond to new **mot_dest** commands.<br>3 = Motor on but power saving (offtime reached, usually low torque, steppers have no current, servos no pulses sent).<br>4 = Motor on<br>>4 Various motor moving states reported by the firmware eg. accel, full speed, decel (in flux in code...) |
| mot_dest | m1-4 [new_destination] | 4.5 | Get/set the current motor destination. This command is non-blocking, the command does not wait for the motor to arrive. Use the mot_wait command if you need that. The destinations do not buffer, ie. if you set a new destination while it's moving, it will start heading to the newly defined destination (including any deceleration required for direction reversal).<br>eg. mot_dest m2 +3200 #will start moving to position +3200<br>eg. mot_dest m2 r+320 #relative move, add 320 steps to the current destination position<br>eg. mot_dest m3 r-n #relative move, step "nudge" steps in the minus direction ( "r+n" for the other way) |
| mot_pos | m1-4 [redefined pos] | | Outputs the current motor position (can be different from the destination value if it's still moving). Writing a |

| | | | |
|---|---|---|---|
| | | | "redefined pos" (steppers only) will redefine the current position to be the value given (ie. this is for position re-definition or zeroing, the stepper motor will not move).<br>NB. You cannot redefine servo positions in this way. In servos the position value is the same as the output pulse-width, in microseconds. |
| mot_out<br>mot_outn | m1-4<br>[new dest] | 10 | If no new destination value, this returns the motor position value in real-world units rather than step (stepper motors) or microsecond (servo) units. If a new destination is set, this takes the real-world units configured in mot_out_conf. The "mot_outn" version does not include the user-defined units in the output.<br>eg.<br>mot_out m6 20.34 // move the stage Z to 20.34 microns above the slide surface |
| mot_out_conf | m1-4<br>[mult=multiplier]<br>[offs=offset]<br>[units="?"]<br>[decp=n] | | Configure the output of the mot_out command. The offset value is subtracted from the motor position before the multiplier is applied. Steppers should not need an offset if you set your min/max and pos.<br>eg. mot_out_conf m6 mult=0.006095 unit="um" decp=3 // Newport TRA12PP stepper in x4 microstepping mode |
| mot_offtime | m1-4 [time] | | Get / set the motor off-time (milliseconds). If zero the motor will stay on (high torque) until mode set to 0. Otherwise the motor driver will turned off, if static for this amount of time. This can reduce motor heating and possibly some vibration or hum. It may also allow the mechanism to be turned manually. For steppers this can cause loss of stepper synch (step position). In servos, the pulse-width signal will stop (Note that some digital servos may remain in the high-torque state after the pulse signal stops). |
| mot_wait | m1-4 | | (macro-only) Pause the macro till the motor has arrived at destination. |
| mot_end_low<br>mot_end_high | m1-4 [motorpos]<br>[stp=0/1] [rst=0/1]<br>[inv=0/1] | | Define a software low end-stop. Any motor destination set below this value, will be set to this value. You cannot set a low endstop above the current step position.<br>The fourth control line can define a hardware endstop, for step position referencing and preventing movement beyond this limit. On lowering step value, if the limit pulls low, the hardware endstop has been reached. Motor position value is redefined to be equal to the "motorpos" value.<br>**Hardware limit switches (Steppers only)**<br>The fourth line (lim) of each stepper motor interface is an input used for hardware limit switches.<br>**stp=0/1** Specifies that the motor should stop abruptly when the limit line is activated. If stp is set, the motor will reverse direction, stepping at the start rate, until the limit signal deactivates.<br>**rst=0/1**Resets the current motor position to be "motorpos" when the limit line de-activates.<br>**inv=0/1**Typically when the limit switch is active the line reads high, set inv=1 where a reading low means an activated limit switch. |

| | | | |
|---|---|---|---|
| mot_lim | | | Returns the state of the stepper motor limit switches. Bitfields alternating low & high limit switches, starting at motor m1. ie. bit 0 for motor m1 low, bit 1 m1 high. eg. a return value of 16 means that motor m3 low limit switch is active. |
| mot_rate | m1-4 [time] | | Sustained pulse rate when turning at full (max) speed. Defined as the interval between step pulses, so **smaller is faster**. There's a low firmware limit ( 5µus ), but you will probably have electronics or mechanical limits that are slower than this. Too fast and your stepper may skip steps, and lose synch. Tune to taste.<br>mot_rate m4 200us |
| mot_accl | m1-4 [new_accel] | | (Steppers only, Hz/sec$^2$) How fast to accelerate / decelerate the stepper. Will depend the inertia of your system, motor size, and driver current capability. Reduce it if you are getting missed steps, or the motor is getting warmer than you would like. |
| mot_bckl | m1-4 [+/-steps] | | Get/set motor backlash. Correct for slack in your mechanical gearing. |
| mot_nudge | m1-4 | | Get/Set the sizeo of a motor nudge. eg. used with keypad "+" and "-" buttons. |
| mot_present | | | (read only) Bit fields showing which motors are configured. Mainly useful for constructing interfaces. |

**Analogue out control (eg. laser power, LED current, stage position)**

The power board (board reference "p") has 8 analogue out channels split into 2 groups.

**Channels ps, pt, pu & pv (0 to 65535 -> 0 to 5V)**

The lower 4 channels are configured for 0 to 5V output. Each channel can be configured for one of the following:

- DAC off (0V output)
- 0 to 5V voltage out eg. for laser power control
- 0 to 1200mA current out control for LED or direct laser diode drive (needs optional module RDC-24-1.2 installed on the board)

**Channels pw, px, py & pz (0 to 65535 -> 0 to 10V)**

Typically configured with 0 to 10V output using an amplifier set to 2x gain. These may be used for fast stage drive control (PI, Madcity etc). To drive an XYZ stage, connect to channels X, Y, and Z if you want the direction keypad keys to work sensibly.

**Linked on/off control TTL**

If selected on digital i/o config page, each analog-out channel can be linked to a corresponding digital line letter for on/off control. For the lower four channels (S, T, U & V) these lines are wired directly to the power board and have an n-channel MOSFET at the analog output stage. This allows faster "ON" and "OFF" control than can be provided by the DAC.

| Command | Arguments | t (µs) | Description |
|---|---|---|---|
| dac_ref | line [newname] | 3.7 | Get/set the reference name for this line. 11 characters max.<br>eg. dac_refr ps laser561 |

| | | | |
|---|---|---|---|
| dac_mode | line [newname] | | Get/set the DAC mode:<br>0 = Off, stay at 0V output.<br>1 = Off, floating (not working yet...).<br>2 = Voltage output.<br>3 = LED driver (RCD-24 module).<br>5 = Stage axis mode.<br>eg. dac_mode ps 3 // LED driver on DAC channel S |
| dac_dest | line [0-65535]<br><br>or<br><br>line [r+/-/*diff] | 11 | Sets the DAC output destination value (16bits, 0 to 65535). Full scale (65535) gives a DAC output of 5V. A relative incremental / decremental / multiplier change can be produced using "r+", "r-" or "r*". If no value given, the current destination value is returned.<br>In a macro, this command is non-blocking, meaning if slew-rate limiting is applied on this channel (see dac_rateL), then the output signal may continue to change over the following commands. If you need macro progress to wait till it arrives, use the dac_wait command.<br>Examples:<br>dac_dest ps 32768 #(channel s, set to half scale (2.5V) out<br>dac_dest pt r+200 #(channel t add 200 to the current destination DAC value<br>dac_dest pz r+n #(channel z add "nudge" units the the current output<br>dac_dest pu r*1.05 #(channel z increase the output by 5%<br>dac_dest pz #(get the value on channel Z) |
| dac_val | line | 2.5 | (read only) Returns the present dac output value. This can be different from dac_dest value if the output is changing with rate limiting active (ramps applied). |
| dac_rate | line [0-32767] | | If rate > 0, this will apply a slew (ramp) rate limit to the DAC output. Set to zero to remove any rate limiting. The value sets the maximum DAC unit change that happens per 100µs write cycle. This can cause dac output changes to span the commands that follow a dac_dest change. Use dac_wait if you need to pause the macro till the dac output change finishes.<br>dac_rate pz 0 # no ramp rate limiting on channel Z<br>dac_rate py 2000 # full scale change would take about 3ms (ie. 0.1ms x 65536 / 2000)<br>dac_rate px 1 # slow ramping. full scale change would take ~ 3.3s (ie. 0.1ms x 65536) |
| dac_wait | [line] | ~ | (macro-only) When slew rate limiting, this will halt macro progress until the DAC channel has reached it's destination (or reached min or max limits). If rate limiting is not applied (eg. dac_rate ps 0), this command does nothing. With no line specified (dac_wait) macro progress will halt until all 8 channels are at their destination value. |
| dac_last | [line] | 3.5 | (read only) Return the time (microseconds) since the last change on the line specified (or most recently changed channel if non-specified). If a rate-limited move is in progress, this function returns zero. |
| dac_min | line [0-65534] | | Set the minimum output value. Usually used with the RCD24-1.2 programmable current module only. Get/set DAC output value where the LED driver current output is zero milliamps. |

| | | | |
|---|---|---|---|
| dac_max | line [1-65535] | | Get/set the DAC output limit when the channel is in voltage out mode. 0 = 0V, 65535 = 5V |
| dac_out | line | 10 | (read only) Returns the channel output value in real-world units rather than DAC units. The formatting of this output can be tuned using the commands **dac_out_conf**. The output of the dac_outn variant does not include the units. |
| dac_out_conf | line [mult=multiplier] [offs=offset] [units="?"] [decp=n] | | Configure the output of the dac_out command. The offset value is subtracted from the dac output value before the multiplier is applied.<br>eg. dac_out_conf ps mult=0.022186 unit="mA" decp=3 // 1200mA LED driver on channel S |
| dac_nudge | line | | Get/set the nudge value. Mostly influences the speed of the change keys on the keypad. |

**Stage control interface**

Configure any combination of motors (mot) and analogue out (dac) for a stage interface. For example you could have a combination of stepper motor XY (mot interface) and piezo Z (dac interface) configured to control your stage XYZ.

| Command | Arguments | t (µs) | Description |
|---|---|---|---|
| stg_conf_x<br>stg_conf_y<br>stg_conf_z | int=interface<br>[invert] | | Configure each axis with a control interface. Include "Invert" to reverse the interface directionality (web page and keypad). Output also shows min/max and unit information. |
| stg_out_x<br>stg_out_y<br>stg_out_z | [r+] newpos | | Get/set the stage position. Move to an absolute position or use "r+0.100" format for a relative change. |
| stg_val_x<br>stg_val_y<br>stg_val_z | | | (read only) Current value in digital units |
| stg_dest_x<br>stg_dest_y<br>stg_dest_z | | | (read only)Destination in digital units (steps, dac values or microsecs if PWM) |
| stg_min_x<br>stg_min_y<br>stg_min_z<br>stg_max_x<br>stg_max_y<br>stg_max_z | | | (read only)Stage min/max values in digital units. |

**Serial communications (SPI and RS232)**

Two types of interface are supported: Serial peripheral interphase, sometimes called the 4-wire bus or SPI and RS232. WOSM boards have 3x SPI connectors which allow data transfer to daughterboards or other controllers that can talk SPI.
Channel designations as follows:
A (SPI) (26pin, slave mode) Is Raspberry Pi pinout compatible. Has 1 slave select line, and 2 spare TTL. (work in progress)
B (SPI) (10pin, master mode) Has 4 CS lines.
C (SPI) (10pin, master mode)
D (RS232, (3pin GND, TX, RX) Low voltage (0V / 3.3V)
E (RS232) (3pin GND, TX, RX)
F (RS232) (3pin GND, TX, RX)

| Command | Arguments | t (µs) | Description |
|---|---|---|---|
| ser_mode | channel<br>modenum | | Get/set the serial comms mode.<br>The applicable mode numbers are:<br>0 Macro comms only (default mode)<br>1 Laser trap daughterboard (channel b only)<br>100 Slave terminal (channel a only, eg. Raspberry Pi comms)<br>eg.<br>ser_mode a 10 # runs a command/response terminal slave on channel b, macro independent |
| ser_config | channel<br>[config register] | | Get/set the serial configuration register (hexadecimal value). For full configuration details see the microcontroller chip or family PDF. Supports 32-bit hex value or alternatively and some key words.<br><br>On the SPI channles (A, B & C) this writes to the SPIxCON register.<br>Legible switches include:<br>**on** or **off** to enable or disable the SPI module (CON bit 15)<br>**cke=0** or **cke=1** Clock edge<br>**ckp=0** or **ckp=1** Clock polarity<br>eg.<br>ser_config a on cke=1 ckp=0 #turn on SPI bus A<br>ser_config a 0x18260 #enable SPI bus A in master mode<br><br>On the RS232 channels (D, E & F) this writes to the UxMODE register.<br>More legible switches include:<br>**on** or **off** to enable or disable the serial module<br>**pdsel=?** data/parity selection: 0 -> 8/none, 1 -> 8/even, 2->8/odd, 3->9/none<br>**stsel=?** stop bits (0 for 1-stop, 1 for 2stop bits)<br>eg.<br>ser_config d on pdsel=2 stsel=0 #turn on serial bus D, 8 data bits, even parity, 1 stop bits |
| ser_brg | channel<br>[divisor] | | Get/set the SPI baud rate generator divisor. Outputs the current divisor value and the resulting bus clock speed in Hz ie. bit-rate, not byte-rate.<br>eg.<br>ser_brg a 0 #set SPI bus A to 20MHz |
| ser_cs | channel<br>[CS line]<br>[low] | | (SPI master mode only) Get/set the SPI chip select line. SPI connectors have up to 4 ( 1 to 4) chip select lines. When a line is selected, all subsequent calls to ser_send will pull this line low when data is transmitted. The line is pulled high at the end of transmission.<br>On call this command will set the line high (usually means "unselected"), unless you add the "low" parameter. eg.<br>ser_cs a 1 #SPI bus A to use CS line 1. This line set high. It will pull low automatically when the wosm sends data<br>ser_cs c 2 low #SPI bus C to use CS line 2, pull it low immediately<br>ser_cs b 0 #SPI bus B will not use any chip select line |
| ser_send | channel<br>data | | Send the data down the bus. The sent data is escaped ascii in double quotes.<br>The following escape characters are supported: \r \n \t \0 \" \' \\ as well as \xFE to define individual hexadecimal characters. eg.<br>ser_send b "reset\r\n"<br>ser_send b "reset\x0D\x0A" |

| | | | |
|---|---|---|---|
| ser_send_rep | channel repeats char | | Send same repeated character down the bus.<br>Max repeat is eg.<br>ser_send_rep b 64 "\xFF" |
| ser_recv | channel [max=?] [ignore=?] | | Recieve data from the SPI channel buffer.<br>The SPI bus master may need to clock out dummy bytes in order to fill this buffer.<br>eg.<br>ser_send b "\xFF\xFF\xFF\xFF\xFF\xFF\xFF\xFF\xFF\xFF\xFF\xFF\xFF\xFF\xFF\xFF" #dummy bytes to read data from device<br>ser_send b "\xFF\xFF\xFF\xFF\xFF\xFF\xFF\xFF\xFF\xFF\xFF\xFF\xFF\xFF\xFF\xFF" ${fsize} = ser_recv b max=32 eol=" " ignore="\xFF" |
| ser_recv_clr | channel | | Clears any data in the receive buffer. |
| ser_wait | channel | | (macro only) Wait for serial hardware to be idle.<br>In SPI bus master mode macro execution blocked until the most recent send is complete.<br>In SPI slave mode the macro will halt here until the SS line is high (the bus master has stopped sending). |
| ser_file_rd | channel filename.ext | (slow) | (2019 still in dev...) Read a file from a daughterboard.<br>The local (WOSM SD card) default folder is /"//wosm//data/".<br>The daughterboard folder is the root folder.<br>First line returned (ascii) contains these details (MD5 included if the "filename.ext.MD5" file exists: filename.ext numbytes fatdate [MD5]\n<br>[binary file data .....] eg.<br>ser_file_rd b newdata.dat |

**Network and TCP/IP**

These commands are used if you're setting a static network config, otherwise the network settings are obtained automatically using DHCP (make sure you have "lan_conf_ip 2" in your config.wml on the SD card. You can see the network settings on the Network page on the keypad.
If you can connect you can make static settings changes on the web gui conf/network page. Else you will need to edit the "wosm/macros/config.wml" file on the SD card directly. Example static network settings below
lan_conf_ip 1
lan_ip 192.168.77.44
lan_subn 255.255.255.0
lan_gateway 192.168.77.1
lan_dns1 192.168.77.1

| Command | Arguments | Description |
|---|---|---|
| lan_host | up to 15 char. | Get/set the computer hostname. |
| lan_conf_ip | 0, 1 or 2 | 0=private address, 1=static, 2=DHCP(default) |
| lan_subnet | | Get/set the lan subnet |
| lan_subn_pfx | | Get the lan subnet as prefix bit count. eg. 24 |
| lan_domain | | |
| lan_ip | | Get/set the lan IP address. To set the address, the lan_config must be set to 1 (static). |
| lan_mac | | Get lan MAC address. |
| lan_gateway | | Get/set the gateway IP. (lan_config must be 1 to set this). |

| | | |
|---|---|---|
| lan_dns1<br>lan_dns2 | | Get/set the lan nameserver IPs. Set requires that lan_config=1. |
| lan_ttl_user<br>lan_ttl_drvr | | You can reduce the TTL of IP packets leaving the controller. Time to live (TTL) defines the IP packet lifetime in terms of network hops. If you connect to a controller across two network switches, your TTL must be more than 2 (TTL >= 3) or the connection will fail due to packet loss.<br>Mainly useful for security. This is simple network firmware. As yet there is no packet encryption (no HTTPS or SSH). A low TTL can rule out internet connections as well as longer range lan connections, independent of any firewall config you may have.<br>You could also use it to count hops to your controller, but you would be better off using traceroute (which also works by incrementing packet TTL) from your host PC. User connection TTL (lan_ttl_user) sets the maximum number of hops for HTTP (port 80) and user Telnet (port 23) connections. Driver TTL (lan_ttl_drvr) is for controller PC comms (Telnet over port 1023).<br><br>NB. On this page you will see two meanings of TTL. It's just an acronym coincidence, they shouldn't be confused. The time-to-live of network packets is rarely used in the same context as transistor-transistor logic, referring to digital signalling between ICs. |

**Macro control commands**

Macros are stored in the folder /wosm/macro/ on the µSD card. You can edit them from there or more usefully via the web gui. Up to 8 macros can be loaded and running simultaneously on the wosm. On first run macros need to be loaded from the micro-SD card, this takes about 7ms on my development system but will depend on your SD card speed and macro size. Subsequent calls to the macro will be much faster (a few microseconds, not milliseconds) as the macro stays in ram. If all macro slots are full, non-running macros are removed on a 'last-used' basis when a new macro needs to be loaded. The functions below do not need the ".wml" extension part of the macro filename.

| Command | Arguments | t (µs) | Description |
|---|---|---|---|
| wml_run<br><br>wml_run_wait | macroname<br>[param1=value]<br>...<br>[param5=string] | | Run the named macro. If called from a macro, both macros will continue simultaneously.<br>If you want the calling macro to wait until the called macro finishes, you should use the **wml_run_wait** version. Optional parameters will be available as variables in the macro.<br>eg.<br>wml_run timelapse2D nframes=1000 expos=100ms intervl=2000ms<br>Will run the macro "timelapse2D.wml". Within this macro commands can refer to the variables "${nframes}" and "${expos}". eg.<br>loop count=${nframes} dur=${intervl} {<br> dig_hilo n ${expos} # camera trigger on line "n"<br>} |
| | macroname | | |
| wml_running | | | (read only) Returns a space-separated list of running macros. |

| | | | |
|---|---|---|---|
| wml_stop | macroname | | Stops the macro.<br>Details: On first call the macro is put in "no_loop" mode. Any loops do not repeat, so usually the macro finishes quite quickly.<br>This is intentional behaviour, that will allow any cleanup or housekeeping commands a the end of the macro to run.<br>If this is called when the macro is already in "no_loop" mode, the macro will be halted abruptly. |
| wml_pause | macroname | | Pause execution of the macro. ( Not functional, Needs work/testing, timed loops will not behave nicely yet..., NJC). |
| wml_unload | [macroname] | | Remove this macro from RAM. If no macro named, then it will try to remove all loaded macros. Running or paused macros are not unloaded. |
| wml_file_new | macroname | | New macro file. |
| wml_file_del | macroname | | Delete this macro from the SD card. |
| wml_file_cat | [number] | | Catalogue / list the defined macros (alphabetical?) on the memory card, returns nothing when number > last file number. |

**Macro flow control**

These commands will only work in WOSM macros. They will not be recognised in any other context, they'll just raise an "unknown" error.

| Command | Arguments | t (µs) | Description |
|---|---|---|---|
| pause | delay_time | | Pause the macro for delay_time. The delay is entered in the wosm time format. The minimum period possible is 15µs on last scope tests (if running "fast", see below). Timing accuracy is about a microsecond. Pause commands less than 30µs are blocking commands, ie. they will not service any other macros or devices during the pause.<br>Also note that a pause longer than 30µs will remove any fast running condition. eg.<br>pause 127us<br>pause 48ms<br>pause 20min |
| loop | [count=?]<br>[dur=?]<br>{...} |  | Loop the curly bracketed commands "count" times. You can nest loops up to 8 levels.<br><br>Use "dur=" to specify a timed loop, in wosm time format. The timing delay for dur occurs during the jump back to the start of the loop ie. the first pass through the loop happens without any delay. and no delay is added to the end of the last pass through the loop.<br><br>Example:<br>loop count=1024 dur=250ms<br>{<br> other<br> commands<br> here....<br>} |

| | | | |
|---|---|---|---|
| | | | This example would loop 1024 times at 4 loops/second (ie. 250ms loop duration).<br>There is an implicit "fast 1" at the start '{' and end '}' of each loop, mostly to give you the chance ot set fast mode in the first command of the loop if you need it. This may be useful for timing-critical start of loop commands. |
| loop_idx | | | Get the interation number at this loop level.<br>loop count=10 dur=1s<br>{<br> ${lp} = loop_idx; #get the loop index<br> kyp_lcd_text x=0 y=0 "Loop index:\${lp}" #show it on the lcd<br>} |
| if | ( v1 < v2 )<br>or<br>( v1 = v2 )<br>or<br>( v1 > v2 )<br>or<br>( v1 != v2 ) | | Fairly basic conditional execution. Values can be numbers or variables or a combination. Values are converted to 64bit floating point numbers for comparison.<br>It's fussy, it will fail if there is not opening and closing brackets (), and the opening curly bracket "{" on the same macro line.<br>As yet no "else" option.<br>Example:<br>${temp} = temp_deg 0<br>${msg} = "Normal"<br>if ( ${temp} > 30.0 ){<br> ${msg} = "HIGH"<br>}<br>if ( ${temp} < 15.0 ){<br> ${msg} = "LOW"<br>}<br>kyp_lcd_text "Temperature:\${temp} C \${msg} " |
| fast | [n] | 0.5 | Run the next 'n' commands as fast as the microcontroller can interpret them. Just use this where you need it, as it blocks other macros and processes.<br>If you omit the 'n' value, the internal fast command count-down counter is set to $2^{32}$. In most circumstances one of the commands below will bring the macro out of fast mode before this counter reaches zero.<br>The following commands bring the macro out of 'fast' mode:<br>**fast 0**<br>**pause**<br>**loop {...}** (really this sets the fast count-down count to 1, see the loop command) |
| stop_on | [conditions] | | By default macros will halt on any error or timeout. There may be conditions where you want the macro to continue to the next command if there is a timeout. The following commands would allow that, for just one dig_wait command.<br>stop_on -timeout # don't stop the macro if a timeout happens<br>dig_wait a 1 1s # wait 1s for line "a" to go high<br>stop_on timeout # any further timeouts will halt as normal...<br>Valid conditions are: all, unknown, timeout<br>Precede conditions with a minus ("-") to remove the condition eg. stop_on -all<br>would remove all possible reasons to halt the macro, probably a bad idea.<br>All macros start with an implied "stop_on all" condition, so there's no need to include that command in the macro. |

**WOSM time format**

Some commands accept a time value (eg. "pause 200ms") which needs to be interpreted quickly. The following restrictions apply to these time values:

- Integer numbers only
- Accepted units are: us, μs, ms, s, sec, min, hr
- Defaults to microseconds (us) if no units specified.

**Macro variables**

Each macro has a limited number of local variables. These are text based.
Variables can be set within the macro, or on the macro call line.
eg.

wml_run acquire2D nframes=2000 expos=200ms
This example would set two variables that can be used in the macro "acquire2D.wml", accessible in the macro as as ${nframes} and ${expos}.

Notes:
* Variable name length can be up to 7 characters
* Variable content can be up to 32 characters
* Macros can have up to 32 local variables
* Local variables do not persist between runs of the macro

**Global variables**

Up to 32 global variables are available to all running macros. They persist after a macro exits. Not wiped till you reboot. Define global variables by prefixing the variable name with "g_".
eg.

${g_count} = ical ${g_count} + 1  #increment a global counter

| Command | Arguments | t (μs) | Description |
|---|---|---|---|
| ${varname} = wosm_cmd | | | To set the variable called "varname" (case sensitive) to the output of the wosm_command. Variable names can be up to 7 characters, variable content currently limited to 16 characters.<br>eg. ${roomtmp} = temp_deg2 |
| ${varname} = "string" | | | To set the variable called "varname" to the string shown. Quotes required, stored as strings.<br>eg.<br>${expos} = "100ms" #set variable 'expos' = '100ms'<br>dig_hilo_t ${expos} #put out a 100ms pulse on line 't' |
| ical<br>fcal | ${v1}<br>operation | 6-<br>15 | Simple calculations.<br>ical works on signed 64bit integers<br>fcal, for floating point calculations (long double, 64bit float).<br>arithmetic operations (ical and fcal): + - / * |

| | | | |
|---|---|---|---|
| | ${v2}<br>["*format*"] | | logical operations (ical only): & (and), \| (or)<br>Calculations must have two values, or variables, and a single operator, more complex operations should split over several lines.<br>The optional format, in quotes, allows you to format the output value: limit decimal places, or write the output in hex etc. (see C printf for more formatting options)<br><br>Examples:<br>${t1} = ical ${t1} - ${t2} # subtract t2 from t1<br><br>${v1} = fcal ${v1} / 10.24 "%.3Lf" # divide v1 by 10.24, save result in v1<br> # (output formatted to 3 decimal places)<br><br>ical (integer) format examples:<br>"%lld" (default) 64bit signed decimal, fmt not necessary<br>"%012lld" 12 character, 64bit decimal integer, with zero padding if necessary<br>"%llx" 64bit hexadecimal eg. "F4E1D21F45ED1"<br>"0x%016llx" 64bit hexadecimal, 16characters including zero padding, '0x' prefix<br> eg. "0x000014EF1ABCDEF7"<br><br>fcal (floating point) format examples:<br>"%Lf" (default), fmt not necessary<br>"%.3Lf" 3 decimal places<br>"%.3LE" output showing exponent, 3 decimal places |
| fn | function<br>var1<br>[var2]<br>["format"] | | Returns a floating point (64bit) answer. Formatting options the same as fcal.<br><br>Function list with examples:<br>${v} = fn pow 2 16 #power 2^16, (v = 65536)<br>${v} = fn sqrt 16 #square root (v = 4)<br>${v} = fn fabs -7.47 #absolute value, removes minus sign (v=7.47)<br>${v} = fn sin 1 #sine<br>${v} = fn asin 1 #arcsine<br>${v} = fn cos 1 #cosine<br>${v} = fn acos 1 #arccos<br>${v} = fn tan 1 #tangent<br>${v} = fn atan 1 #arc tan<br>${v} = fn ln 2 #natural log, (v = 0.6931471805599)<br>${v} = fn exp 0.69314718 #exponential, (v = 2) |

**System versions and updates**

Here for my reference (NJC). Not too useful for macros.

| Command | Arguments | t (μs) | Description |
|---|---|---|---|
| sys_file_app<br>sys_file_boot | | | Read only. Returns the name of the file flashed on the last firmware update, either for the main wosm firmware (app) or the bootloader (boot) |

| | | | |
|---|---|---|---|
| sys_fdat_app<br>sys_fdat_boot | | | Read only. Returns the creation date of the file of the last firmware update, main app or bootloader. |
| sys_ver_app<br>sys_ver_boot | | | Read only. Returns the app or bootloader firmware version string. |
| sys_sdfw_app<br>sys_sdfw_boot | | | Read only. Returns the filename (without the .hex extension) of the most recent 'flashable' update present on the sd card. This is used to populate the update web page. I can't think of another possible reason you would want to use it.<br>In order to be listed:<br>* File must be dated (created) later than the current firmware.<br>* App updates must be named mz_??????.hex, bootloader updates named mzbl????.hex |
| sys_flash_app<br>sys_flash_boot | 8char | | Perform a wosm app or bootloader firmware update. The app requires a reboot/reset to complete on next wosm bootup. The firmware hex image must be on the SD card, in the folder "/wosm/firmware/". The file must have the ".hex" file extension, and the filename cannot be more than 8 characters. App firmware must be named MZ_?????.hex, bootloarder firmware is called MZBL????.hex. The ????? is usually the firmware version number.<br>eg. sys_flash_app MZ_0912<br>will flash the hex file "/wosm/firmware/MZ_0912.hex" on next wosm reset or boot. It takes a few seconds to complete. Returns the name of the pending flash file....?<br>Notes:<br>* the word "latest" will flash the most recent hex in the firmware folder.<br>* cutting the power while flashing is in progress is deemed unfriendly.<br>* This command will not work in a macro (command-line or web click only)<br>* the firmware flash performs a crc check on the entire file before any flash happens.<br>* The bootloader can only read cards that are FAT32 formatted (although wosm firmware can work with other formats) |
| sys_board | | | Returns "MX" or "MZ" depending on microcontroller type. |
| sys_upd_val | | 20 | Time (unixtime) of the most recent update check (SD card files). |
| sys_upd_str | | 37 | When was the most recent online HTML page update (SD card files). |
| sys_upd_mode | | | Automatic web updates. 0=Off, 1=Every boot, 2=Weekly (within 2min of boot) |
| sys_upd_now | | | Update the WOSM control pages from wosmic.org. Runs in the background, will take a minute or so to complete. New web pages are functional immediately. New firmware (hex) images may also be downloaded to the SD card during this update process, new firmware images will not be flashed automatically, you must select firmare updates on the update config page. |

**User management**

Users login at HTTP and Telnet connections. The default user "admin" cannot be deleted. At the moment user login is simply for network login.

| Command | Arguments | t (µs) | Description |
|---|---|---|---|
| usr_set | username [MD5=passwordMD5] or username [pass=password] | | If the user does not exist, is created. Also sets the password for this user:<br>usr_set barry pass=mysecret<br>Note that only the password hash (MD5) is saved. |
| usr_del | username | | Remove this user. You cannot remove the default "admin" user. This does not delete user macros from the SD card. |
| usr_name | [0-15] | | If no arguments, returns the currently authenticated user. The number argument returns the user name from the user database. |

**Data buffer commands**

Work in progress. These commands work with the Wosm64 data acquisition software (not with Micromanager at this stage). Mostly concerned with getting a low bandwidth image to the browser, for remote monitoring acquisition progress.

| Command | Arguments | t (µs) | Description |
|---|---|---|---|
| dat_name | | | (read only) Read the the (file)name of the data buffer. |
| dat_size | | | (read only) Return the size (bytes) of the data in the buffer. |
| dat_max | [SetMax] | | (read only) Return the maximum buffer data size. |
| dat_recv | name | | [Telnet port 1023 only]. Binary receive the data. |

**HTML commands**

Here for reference only. These commands provide dynamic HTML content for fast web page loads. Not too useful for macros.

| Command | Arguments | t (µs) | Description |
|---|---|---|---|
| htm_sign | | | |
| htm_edt1 | | | |
| htm_edt2 | | | |
| htm_mnu_int1 | | | |
| htm_mnu_int2 | | | |

**Command speed test macro**

Timing information on this page is measured in the script below. The command to be timed is run 10 times in fast mode. With this digital output example you can confirm the timing on a scope. Just make sure that line q is configured to be an output.

```
fast
${tz} = sys_usec
########### 10 commands to time between here...
dig_out q 1
dig_out q 0
dig_out q 1
dig_out q 0
dig_out q 1
dig_out q 0
dig_out q 1
dig_out q 0
dig_out q 1
dig_out q 0
############ ... and here.
${t} = sys_usec
${t} = ical ${t} - ${tz}            #subtract start time of command block
${t} = ical ${t} - 6                #correct for timer read delay
${t} = fcal ${t} / 10  fmt="%.1Lf"     #divide by n=10
kyp_lcd_screen t=1s "Command time:\n\n    ${t} us"
```
