## Supplemental Table 2 for "The eduWOSM: a benchtop advanced microscope for education and research"

Carter et al **Supp. Table 2** eduWOSM Bill of Materials

*Includes alignment tools and equipment but not PC. Prices obtained Spring 2024.*

| *Part* | *PN* | *Cost* | *Total* | *Alternative* | *Cost* | *Total* |
|---|---|---|---|---|---|---|
| **LED Illuminator** | | | | | | |
| Luxeonstar Red | LXZ1-PD01 | $6.90 | | Luxeonstar Quad | $27.00 | |
| Luxeonstar Lime | LXZ1-PX01 | $7.90 | | Thorlabs AC127-030-A | $59.40 | |
| Luxeonstar Blue | LXZ1-PB01 | $6.00 | | Thorlabs Lens tube SM05L05 | $15.25 | |
| Luxeonstar 395 | LHUV-395-A065 | $21.48 | | Thorlabs Lens Tube SM05L10 2x | $32.66 | |
| Knight Optical Blue/UV dichroic | 420 FDL | $53.00 | | | | |
| Knight Optical Blue/green dichroic | 565 FDL | $53.00 | | | | |
| Knight Optical Green/Red dichroic | 635 FDL | $53.00 | | | | |
| Thorlabs 30 mm achromat, 0.5 inch dia | AC127-030-A | $59.40 | $260.68 | | | $134.31 |
| **Dichroic Filters / Mount** | | | | | | |
| Chroma Quad LED | 89402 | $1,550 | | Semrock Quad LED-DA/FI/TR/Cy5-B-000 | $1,920.00 | |
| Thorlabs M3 nylon set screws 10 pk | SSEMN4 | $15.13 | | | $15.13 | |
| Thorlabs 1" square broadband mirror | BBSQ1-E02 | $81.22 | $1,646 | | $81.22 | $2,016.35 |
| **Camera Pathway** | | | | | | |
| Thorlabs 1 inch lens tube 0.5" long | SM1L05 | $13.62 | | | $13.62 | |
| Thorlabs 1 inch lens tube 2.0" long | SM1L20 | $17.85 | | | $17.85 | |
| Thorlabs 1 inch 75 mm achromat | AC254-075-A | $87.35 | | | $87.35 | |
| Thorlabs 1 inch lens tube to C-mount | SM1A9 | $20.96 | | | $20.96 | |
| FLIR Blackfly S | BFS-U3-16S2M-CS | $371 | $510.78 | FLIR Blackfly S BFS-U3-13Y3M-C | $480.00 | $619.78 |
| **Objective** | | | | | | |
| Nikon 100x oil 1.3 NA Plan Fluor lens | MRH 01902 | $2,165.97 | $2,165.97 | | $2,165.97 | $2,165.97 |
| **Block** | | | | | | |
| Wolf River Machine | | $1,120.29 | $1,120.29 | | $1,120.29 | $1,120.29 |
| **Sample Stage** | | | | | | |
| Thorlabs Z-axis | MT1B/M | $255.05 | | | $255.05 | |
| Thorlabs X/Y axis 2x | MT1/M | $680.14 | | | $680.14 | |
| Thorlabs base | MT401/M | $26.36 | | | $26.36 | |
| Thorlabs Right angle bracket | MT402 | $58.82 | | | $58.82 | |
| Thorlabs side mount for actuator 2x | MT405 | $132.22 | | | $132.22 | |
| Neodymium Magnet from Apex Magnets | 6 mm x 2 mm | $10 | | | $10.00 | |
| Newport TRA12PPD Stepper | TA12PPD | $1,002 | $2,164.59 | Sparkfun Stepper ROB-10846 | $19.50 | $1,182.09 |
| **Electronics** | | | | | | |
| Elecrow Est for now | | $750 | $1,000 | | $750 | $1,000 |
| **Accessories** | | | | | | |
| Giottos Rocket Air Blaster Large | AA1900 | $17.99 | | | $17.99 | |
| Thorlabs Wrench for lens tubes | SPW602 | $30.64 | | | $30.64 | |
| Metric Allen Wrench Set Amazon | | $15.00 | | Metric set Thorlabs CCHK/M | $28.12 | |
| iexcell 250 Pcs M3 x 6/8/10/12/16 Alloy Steel 10.9 Grade Hex Socket Flat Head Cap | | $9.68 | | | $9.68 | |
| M4 flat head hex set Amazon Black Oxide or Stainless | | $15.00 | | | $15.00 | |
| M3-M6 socket head set Amazon VIGRUE 304 | | $21.99 | | | $21.99 | |
| M25x0.75 thread - specialty tap Amazon | | $30.00 | $140.30 | Thorlabs | $157.00 | $280.42 |
| **Alignment Optics** | | | | | | |
| Thorlabs 6" 1 inch lens tube | SM1E60 | $52.42 | | | $52.42 | |
| Thorlabs 3" slotted lens tube | SM1L30C | $77.45 | | | $77.45 | |
| Thorlabs Frosted alignment disk 2x | DG10-1500-H1-MD | $75.14 | | | $75.14 | |
| Nikon thread to lens tube adapter | SM1A11 | $23.53 | | | $23.53 | |
| Thorlabs 639 nm USB diode laser | PL202 | $134.12 | | Thorlabs Handheld Diode Laser HLS635 | $761.25 | |
| Thorlabs adapter for 1" lens tube | AD11F | $33.99 | | Thorlabs fiber P1-630A-FC-1 | $78.73 | |
| | | | | Thorlabs F230FC-B Fiber collimation | $177.75 | |
| | | | | Thorlabs AD1109F collimator to 1/2" lens tube | $35.44 | |
| | | | $396.65 | Thorlabs SM1A6 1/2" to 1" lens tube adapter | $22.37 | $1,304.08 |
| **Total** | | | | | | |
| **Cheaper version with stepper** | | | **9405.61** | **Fancier version with stepper** | | **9823.29** |
